# Computational predictions and evolutionary analysis of *LrK10* kinase-related putative *PSTOL1* gene homeologs in wheat and orthologs of its wild relatives

**DOI:** 10.64898/2026.02.12.702618

**Authors:** Karthikeyan Thiyagarajan, Carolina Saint Pierre, Chitranjan Kumar, Doyeli Sanyal, Garima Thakur, Deepika Singh, Deepshikha Thakur, Ajay Tomar, Prashant Vikram, Ravi Valluru

## Abstract

Phosphorus Starvation Tolerance 1 in rice (*OsPSTOL1*, known as *Phosphorus uptake 1, Pup1*) is a receptor-like cytoplasmic protein kinase that confers tolerance to phosphorus deficiency. The *OsPSTOL1* gene possesses a Ser/Thr kinase and shows high amino-acid sequence similarity with the *leaf rust receptor-like kinase* (*OsLrK10*). We hypothesize that the putative wheat *TaPSTOL1* and *TaLrK10* have a common ancestral origin and that putative *TaPSTOL1* diverged recently acquiring new structural modifications and biological functions in the process. In this study, we identified all putative *TaPSTOL1* homeologs and examine the evolutionary relationship between *TaPSTOL1* and *TaLrK10* in Triticum species. Our results indicate that the putative *TaPSTOL1* diverged recently without possessing the amino-terminal domain, which is a typical characteristic of *TaLrK10*. We observed numerous conversions tracts between these two genes and the substitution pattern of randomly selected amino acids indicates that dynamic selection pressures acted on both genes. The putative *TaPSTOL1* shows high nucleotide diversity compared to *TaLrK10* within *Triticum* species. Further, a multiple-sequence analysis reveals that the third exon of *TaLrK10* appears to have been duplicated and diverged as a putative single-exon based *TaPSTOL1* in bread wheat. Overall, our comparative analysis indicates that both *TaPSTOL1* and *TaLrK10* appears to have diverged from a common ancestor, acquiring distinct structural organizations and biological functions.

## Introduction

Phosphorous (P) is an essential macronutrient constituting approximately 0.2% of the total dry weight in plants [1]. It is involved in many vital processes such as photosynthesis [2], lipid metabolism [3], nucleic acid synthesis [4] and signaling [5]. However, due to the low levels of bioavailability and its slow diffusion in the soil (diffusion coefficient of 10^−7^ to 10^−9^ cm^2^ s^−1^), P uptake is a constraint for the plant [6,7]. It is therefore essential to identify and understand the genetic mechanisms regulating the absorption and utilization of P availability.

A genetic locus conferring P-deficiency tolerance (*Phosphorus uptake 1*, *Pup1*) was previously identified in an *aus*-type Indian rice variety, Kasalath [8]. Near-isogenic lines with *Pup1* locus harbouring Kasalath allele showed 3-fold higher P-uptake and grain yield under low-P conditions [8]. Subsequently, a gene encoding Ser/Thr kinase of the Receptor-like Protein Kinase LRK10L-2 subfamily from the *Pup1* locus, *Phosphorus Starvation Tolerance* 1 (*OsPSTOL1*), was identified as the most likely gene responsible for the phenotype [9]. *OsPSTOL1* is a receptor-like cytoplasmic protein kinase (*RLCK*) that show phosphorylation activity. In plants, about 1-3% of the total number of genes encoding for putative proteins showing kinase activity that is involved in many metabolic processes [10]. Further, lines overexpressing *OsPSTOL1* showed approximately 30% higher grain yield in comparison with the null lines [9]. Indeed, studies identified several novel single nucleotide polymorphisms (SNPs) of *OsPSTOl1* in rice [11,12,13] and haplotypes based on SNPs in wild rice *O. rufipogon* are associated with significant differences in root length and root weight [9].

Subsequently, using *OsPSTOL1* amino-acid sequence as a query, several *PSTOL* homologs were identified in several crop species including six *SbPSTOL* homologs in sorghum [14] and six *ZmPSTOL* homologs in maize [15]. These identified proteins enhanced plant performance, grain yield and root system architecture traits under low-P and were clustered together with *OsPSTOL1* from rice. Further, these identified proteins indicate that the sequences were conserved mostly at the kinase domain. In wheat *TaPSTOL1* like gene was resequenced (from chromosome 5A) from 27 different bread wheat and durum wheat accessions, characterized the expression of *TaPSTOL1* under different P concentrations and demonstrated the induction of promoter in root tips and root hairs under low-P conditions [16]. These studies emphasize that the *PSTOL1* gene has the potential for molecular breeding applications to improve crop performance under low-P conditions.

The *OsPSTOL1* protein shows a high amino acid sequence similarity with the *leaf rust receptor-like kinase* (*LrK10*), located in the *Leaf Rust Resistance* (*LRR*) locus that belongs to the family of *Receptor-Like Kinases* (*RLKs*) [9,17]. Further, *TaLrK10* is involved in leaf rust disease resistance in bread wheat [18]. Hence, *PSTOL1* and *LrK10* belong to *RLCKs* and *RLKs*, respectively, and both carry a Ser/Thr kinase phosphorylation domain. *RLKs* typically have an amino-terminal domain protruding into the extracellular space while *RLCKs* lack this extracellular domain [19]. Previously, six copies of *Lrk* genes were identified by Southern hybridization on the three homeologous group 1 chromosome in wheat [20,21]. The evolutionary relationships between all of the members of the *Lrk* gene family from the A, B, and D genomes of wheat and its progenitors suggest that *Lrk* copies in hexaploidy wheat were phylogenetically sisters to the orthologous copies in the parental diploid species suggesting that the sequences of the homeologous *Lrk* genes evolved independently after polyploidization [22]. Interestingly, assessing the paralogous and orthologous relationships among the *Lrk* genes indicated that gene losses have occurred for this gene family in the Triticeae [22]. Two paralogous loci comparisons further highlighted that while *Lrk* gene content and organization are well conserved, some of the noncoding regions showed a great extent of reshuffling, indicating several local duplications, deletions, and insertions. It has been opinioned that small duplications of *Lrk* gene sequences with no repetitive elements have occurred successfully in Triticeae [22].

As both *PSTOL1* and *LrK10* shows high amino-acid sequence similarity with each other [9,17], we hypothesized that the putative *PSTOL1* is derived from *LrK* sequences through duplication which then diverged as a putative *PSTOL1* acquiring a distinct biological functionality during the process. Here, we identified all the putative *TaPSTOL1* homolog sequences in wheat and examine the evolutionary relationship between putative *TaPSTOL1* and *TaLrK10* in *Poaceae* species focusing on the tribes of *Triticeae*, *Brachypoideae*, *Paniceae,* and *Andropogoneae*. Our results support the hypothesis showing that the putative *TaPSTOL1* has a high nucleotide identity (96%) with the third exon of *TaLrK10* in wheat.

## Material and methods

### Putative TaPSTOL1 and TaLrK10 gene sequences from Poaceae members

We initially used the rice *OsPSTOL1* gene sequence (Accession ID: KU922625) [23] as a reference to identify close orthologs in the bread wheat cv. Chinese spring reference genome [24, 25]. Based on this query, we found top hit gene models TraesCS3A02G261800, TraesCS3B02G295000, TraesCS3D02G261800 from group 3 chromosomes of wheat which we further used 3DL ortholog (contig: IWGSC_V3_chr3DL_scaffold_639, derived through rice *OsPSTOL1* using URGI-IWGSC wheat BLASTN server https://wheat-urgi.versailles.inra.fr/Seq-Repository/BLAST) due to nascent polyploidization (AABB+DD) of D genome to identify other homeologs across all chromosomes of bread wheat and named them as putative *TaPSTOL1-like* gene sequences. The presence of the kinase domain in these sequences was then predicted using Pfam Sequence search [26] and Motifscan (https://myhits.isb-sib.ch/cgi-bin/motif_scan). Further, using the 3DL sequence as a reference, we searched for homologous/orthologous sequences from wheat wild progenitors such as *T.urartu*, *A.speltoides*, and *A.tauschii* and members of the tribes of *Triticeae*, *Brachypoideae*, *Paniceae* and *Andropogoneae* using different public databases such as NCBI (https://blast.ncbi.nlm.nih.gov), URGI (https://urgi.versailles.inra.fr), and *Aegilops* (http://aegilops.wheat.ucdavis.edu). Query coverage of sequences with more than 70% identity was retrieved using BLASTN [27] with E-value 0 to 1e^-130^. In total, 108 DNA sequences from different *Poaceae* species were aligned together using ClustalX [28].

### Evolutionary analysis of putative TaPSTOL1 gene sequences

A phylogenetic tree was generated using Neighbor Joining method [29] with 1000 bootstrap resampling [30]. The evolutionary distance was estimated using the Maximum Composite Likelihood method [31]. Disparity index [32] using Monte Carlo test with 500 replicates was estimated using MEGA [33] Gamma statistic for gene flow estimates of haplotypes [34,35], Nst [36], Fst [37], synonymous to non-synonymous substitutions [38] and other population genetics analysis was performed using DNAsP V5 [39]. The genomic to mRNA alignment was performed using SPLIGN [40]. The prediction of MITE (Miniature Inverted Repeat Transposable Element) terminal inverted repeat was performed using EMBOSS inverted repeat server, and target site duplication was manually assessed. The miRNA prediction was performed in miRbase [41] and target genes for the miRNA were predicted using psRNATarget [42]. Promoter motifs prediction was done using PLACE server [43].

### Evolutionary analysis of putative TaPSTOL1 and TaLrK10 protein sequences

Ortholog and paralog sequences of the putative *TaPSTOL1* protein from diverse species belonging to the tribes of *Triticeae*, *Brachypoideae*, *Paniceae,* and *Andropogoneae* (with a query coverage >70%, identity >85% and E-value: 0 to 3e^-150^) were retrieved from NCBI. The sequences were aligned using ClustalX using default parameters [28]. Subsequently, a phylogenetic tree using the Neighbor Joining method [29] with 1000 bootstrap resampling [30] evolutionary distance using Jones-Tailor-Thorns (JTT) method [44], RelTime relative divergence tree [45] and Tajima neutral evolutionary and relative rate test [46], rates of amino acid substitutions using the JTT [47] model through Maximum log-likelihood method, and disparity index [32] were generated in MEGA7 [33].

### Retrieval of protein PSTOL1 and LrK10 gene/protein sequences in wheat and close relatives

For comparative genomic study, an ancestrally kinase related *TaLrK10* gene from *T.aestivum* was retrieved from the NCBI nucleotide database (accession: U51330) [18] and was further subjected to BLASTP to retrieve the related sequences from *T.aestivum*. Further alignments with *TaPSTOL1-3DL* revealed that the *T.aestivum Kinase 2* (*receptor-like kinase 2*; Genbank accession: ACK44484) is closely related to the putative *TaPSTOL1* and was therefore included in phylogenetic and protein sequence analysis. Further, The BLASTP analysis of putative *TaPSTOL1-3DL* revealed the highest amino acid sequence similarity (100% query coverage and 96% identity) with putative *AtLrK10* isoforms X1, X2 and X3 from *A.tauschii*.

### Protein 3D modeling and annotations

Protein 3D modeling was performed using Geno3D [48] and annotation and visualization were carried out using the RASMOL Viewer [49]. Kinase domains and active signatures were predicted using Scanprosite tools, and the documentation of protein physicochemical parameters including instability index was calculated using the Protparam tool using default parameters at Expasy (https://web.expasy.org) [50]. Protein motifs were predicted using Eukaryotic Linear Motif (ELM) resource [51]. Functional similarities with the existing X-ray crystal-based PDB structures (5LVO), gene ontology and superfamily superposition were predicted and annotated using the CATH database [52]. Outlier homologs of known structure, PFAM domains, signal peptides, and internal repeats were predicted using the SMART (Simple Modular Architecture Research) [53] and Lifetree was generated using iTOL [54]. Dot plot representation was done using EMBOSS online server [55].

### Expression of putative TaPSTOL1 homeologous sequences

We obtained the gene expression data of putative *TaPSTOL1* homeologs from wheat expression database (http://www.wheat-expression.com), which was already published [56]. Briefly, a 14-d old Chinese spring seedlings (CS) were subjected to P-starvation for ten-days, and root and shoot samples were collected at 0 days (control) and at ten-days (P-starvation). Total RNA was extracted from all tissues and sequencing was performed on each library to generate 100-bp paired-end reads for transcriptomic sequencing on an Illumina High-Seq 2000 platform [56]. We have downloaded the raw data from wheat expression database and mean expression (Transcripts per Million-TPM) was obtained for all the putative *TaPSTOL1* homeologs.

## Results

### Bread wheat has eight putative TaPSTOL1-like gene sequences

Using rice *OsPSTOL1* amino acid sequence as a query (Accession: KU922625) [23], we identified nine wheat sequences with high similarity to *OsPSTOL1*. We observed the first hit on chromosome 5A (E-value 0.0, Milner et al, 2018) and subsequent hits on group 3 chromosomes (3A, 3B, and 3D with E-value ranging from 2e^-139^ to 7e^-138^). Due to close sequence similarity with *OsPSTOL1* (S1 Fig) and recent natural hybridization of the D genome into hexaploid wheat, we further chosen the 3DL homeolog sequence as a query to search for homologs/orthologs in closely related species and wheat progenitors. We identified eight putative *TaPSTOL1* homeolog sequences in the bread wheat cultivar Chinese spring that share 96% identity at the amino acid level. These homeologs are present on chromosomes 3AL, 3B, 3DL, 5AS, 6DS, 7AL, 7BL, and 7DL with an identity ranging from 68% to 95% (S1 Table). We further identified five putative paralogs in durum wheat with nucleotide identities ranging from 67%-95%. Four putative paralogs were identified in wheat progenitors *T.urartu* (67%-95% identity) and *A.speltoides* (65-95% identity), and two putative paralogs in *A.tauschii* (68-96% identity). Similar to *OsPSTOL1*, all eight putative wheat *TaPSTOL1-like* sequences identified are predicted to have a conserved Ser/Thr kinase domain that are conserved across all phyla (Fig. 1).

**Fig 1.**
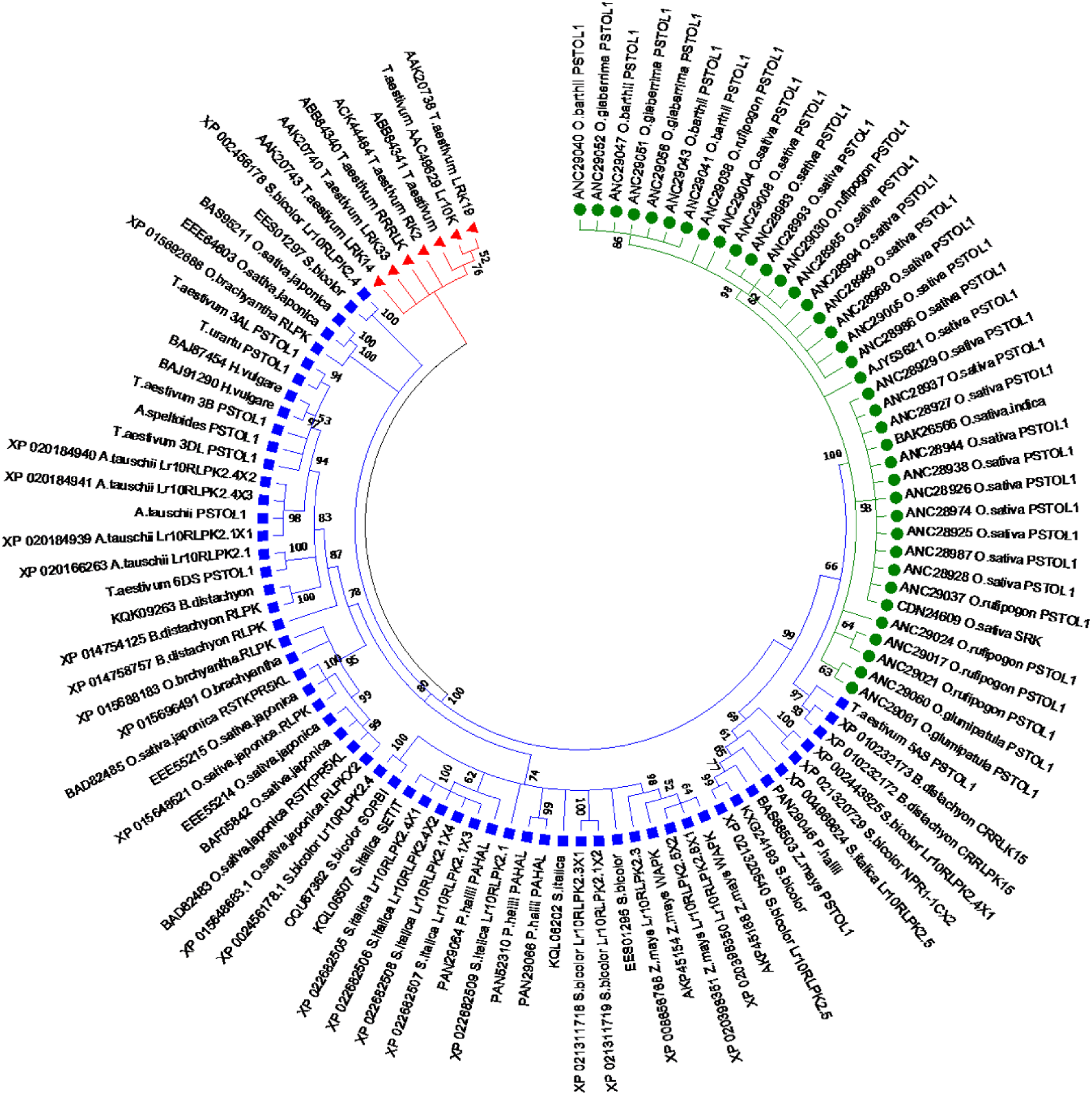
Phylogenetic tree based on the protein sequences of *PSTOL1* and *LrK10* in diverse *Poaceae* members. Clade I: Round green shape symbol: *Oryza* species consists of *PSTOL1* and *receptor like kinase,* Clade II: Blue Square Shape: *Triticeae* and other *poaceae* members with *PSTOL1* and *Receptor kinase* genes, Clade III: Red Triangle shape: *Triticum aestivum receptor like kinases*.

The gene expression profiles of putative *TaPSTOL1-like* were queried in Chinese spring grown under P-starvation for 10 days by probing the wheat expression database (www.wheat-expression.com, queried on April 13^th^, 2019; was already published [56]. In general, all homeologs of putative *TaPSTOL1-like* were more expressed under low-P as compared to the controls (Fig. 2a). A tissue-specific expression pattern was observed for all homeologs in response to 10-d P-starvation (Fig. 2b), suggesting that all the putative *TaPSTOL1-like* homeologs identified in the study are expressed under low-P conditions.

**Fig 2.**
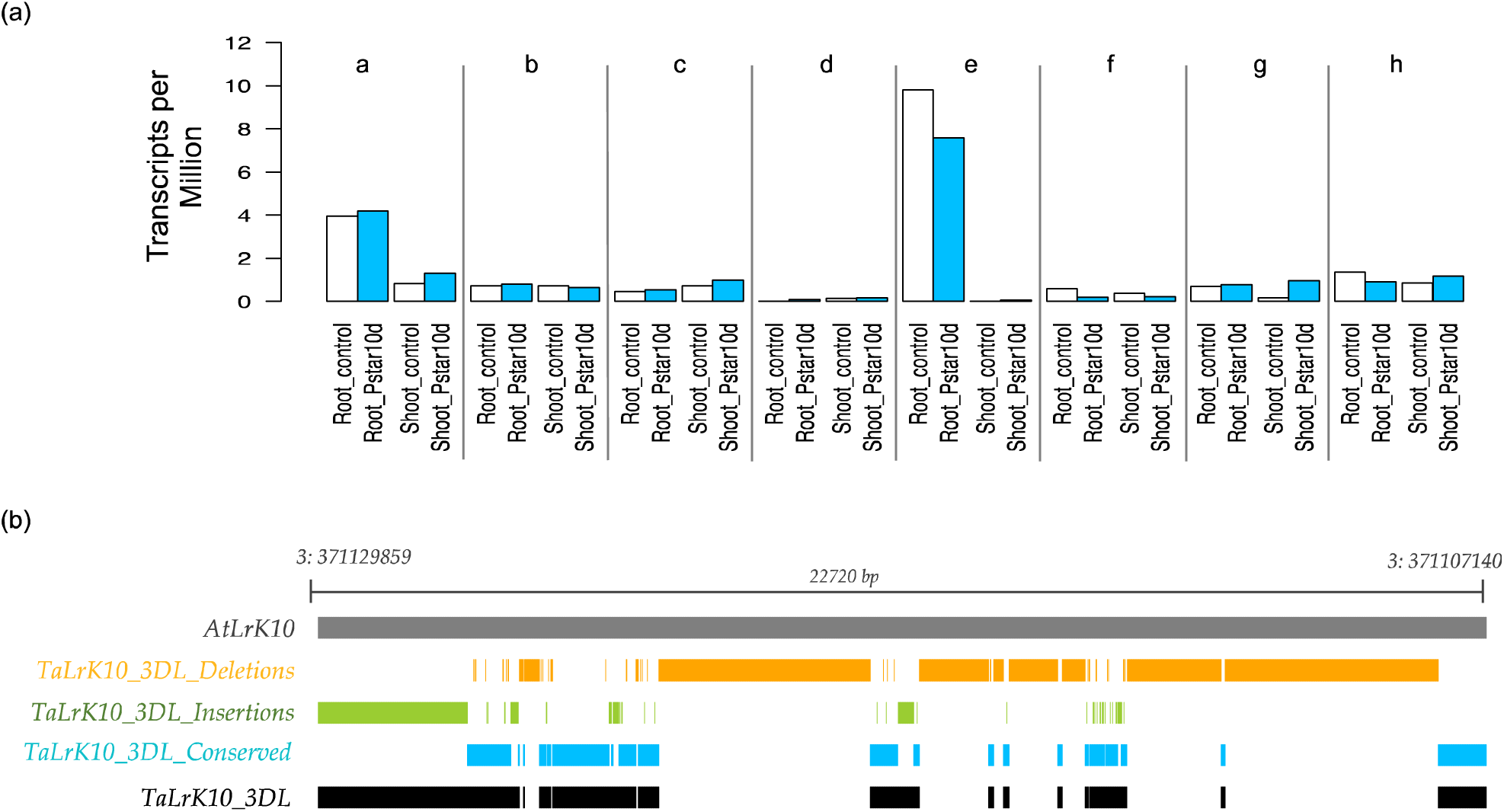
(a) Expression of putative *TaPSTOL1-like* homeologs grown under control (0 days) and P-starvation (10 days) in Chinese spring (Oono et al, 2013; a-h: *Traes_3AL_B27A64367*, *Traes_3B_34896ED29*, *Traes_3DL_E468838BA*, *Traes_5AS_841ECFF18*, *Traes_6DS_84B6888*, *Traes_7AL_F046FDB9E*, *Traes_7BL_504698D18*, and *Traes_7DL_E183F108C*, respectively). (b) Comparison of nucleotide differences between putative *AtLrK10* and *TaLrK10*. Considering the At*LrK10* as an ancestral gene, we identified and plotted both INDELs and conserved nucleotides in *TaLrK10_3DL* in bread wheat. *At, Aegilops tauschii*; *Ta, Triticum aestivum*; 3DL, chromosome 3DL; Pstar10d, P-starvation for 10 days.

We further identified putative *LrK10* sequences in bread wheat and its wild relatives using BLASTN (in NCBI) search with the option MEGABLAST. This search revealed a gene model (*Lr10*, *LOC109770626;* hereafter as *AtLrK10;* Genbank accession for *mRNA*: XM_020329350) that likely showing alternative splicing of three transcript variants whereby X1 and X2 have three exons while transcript variant X3 has four exons (S2a Fig.). The second exon (from transcript variant X3) appears to have lacked in transcript X2 and X1. The third and fourth exons shared the same genomic positions; however, the first exon genomic position vary between three transcript variants (S2a Fig). The presence of *TaLrK10* from bread wheat was previously reported on chromosome 1A (Feuillet et al, 1997). We identified a gene model from IWGSC (*TraesCS3D02G261800*) with exon/intron boundaries harboring four transcripts (hereafter denoted as *TaLrK10_3DL*). This gene locus harbors four transcripts, among which transcripts one and three appear to have protein-coding carrying three exons each while transcripts two and four appear to have nonfunctional (S2a Fig). Overall, the BLASTN searches identified several putative homeologs, paralogs, and orthologs of *TaPSTOL1-like* and *TaLrK10* sequences in wheat and its wild relatives. Further, all putative *TaPSTOL1-like* homeologs expressed more under low-P and were predicted to have numerous transcription factor binding motifs in their upstream region (−1200 bp) of the transcript start site associated with the regulation of RLKs (S2b Fig).

### The third exon of TaLrK10 appears to have duplicated as putative TaPSTOL1-like that carry a MITE insertion which is not present in TaLrK10

To understand the identity between *AtLrK10* and *TaLrK10_3DL*, we further compared both *AtLrK10* and *TaLrK10_3DL*, which revealed that the putative *TaLrK10_3DL* had a 95% identity with the transcript X1 of *AtLrK10*, while the third exon of *AtLrK10* showed an identity of 96% (S3 Fig). Indeed, many INDELs (Insertions: 19% and deletions: 71% relative to the *AtLrK10*) were identified in *TaLrK10_3DL* while 24% of nucleotides appear to have conserved between *AtLrK10* and *TaLrK10_3DL*. We also compared the putative *TaLrK10_3DL* with the *TaLrK10* gene from chromosome 1A [18], indicating that the *TaLrK10* from chromosome 1A shared 58.49% sequence similarity with *TaLrK10_3DL* and 57.09% sequence similarity with *TaLrK10_3AL*.

We hypothesized that the putative *TaPSTOL1-like* is duplicated and diverged from *TaLrK10*. To understand the commonality between them, we aligned sequences of *TaLrK10_3DL* and putative *TaPSTOL1-3DL* from bread wheat (S3 Fig). We noticed that the physical genomic location of *TaLrK10* from both *A.tauschii* and bread wheat overlaps with the putative *TaPSTOL1-like* sequence, suggesting that both sequences are present within the same locus region. Our computational analysis further revealed that the third exonic region of *TaLrK10* (with reference to homeolog from 3D) appears to have duplicated and diverged as the putative *TaPSTOL1-like* gene. Indeed, the third exonic region of *AtLrK10*, *TaLrK10_3DL,* and putative *TaPSTOL1-3DL* shared an identity of 96% (S3 Fig). The other two exons encoding the extracellular N terminal domain in *TaLrK10* did not show any sequence similarity with *TaPSTOL1-3DL* due to the absence of N terminal domain in putative *TaPSTOL1-3DL*. Overall, our analysis suggests that the third exon of *TaLrK10* (location in 3DL: 363607922 bp to 363608939 bp) could indeed act as the putative single exon-based *TaPSTOL1* gene.

We further observed an insertion of MITE (Miniature Inverted Repeat Transposable Element) in the 5′UTR of putative *TaPSTOL1-like* from chromosome 3AL in bread wheat (approx. 90bp in length, containing a 42bp TIR (Terminal Inverted Repeat) and a 2bp Target Site Duplication (TSD)). This MITE is absent in the 5′UTR region of *TaLrK10* from chromosome 1A, while it has many mutations in putative *TaPSTOL1-3DL* and *TaLrK10_3DL* in bread wheat. The identified MITE encodes for the miRNA tae-MIR1137b MI0030384 (Fig. 3), which was previously reported to have highly expressed in mature embryos of wheat during grain development [57,58]. Among wheat progenitors, this MITE is only present in *T.urartu* (S2 Table). In tetraploid durum wheat, the available information on only one of the two homeologous copies of this gene suggests that the MITE insertion is present only on the chromosome 3A while presumably absent on the chromosome 3B. Consistently, the MITE insertion was observed in the putative *TaPSTOL1* on chromosomes 3A and 3D (with many mutations) but absent on chromosome 3B in bread wheat.

**Fig. 3.**
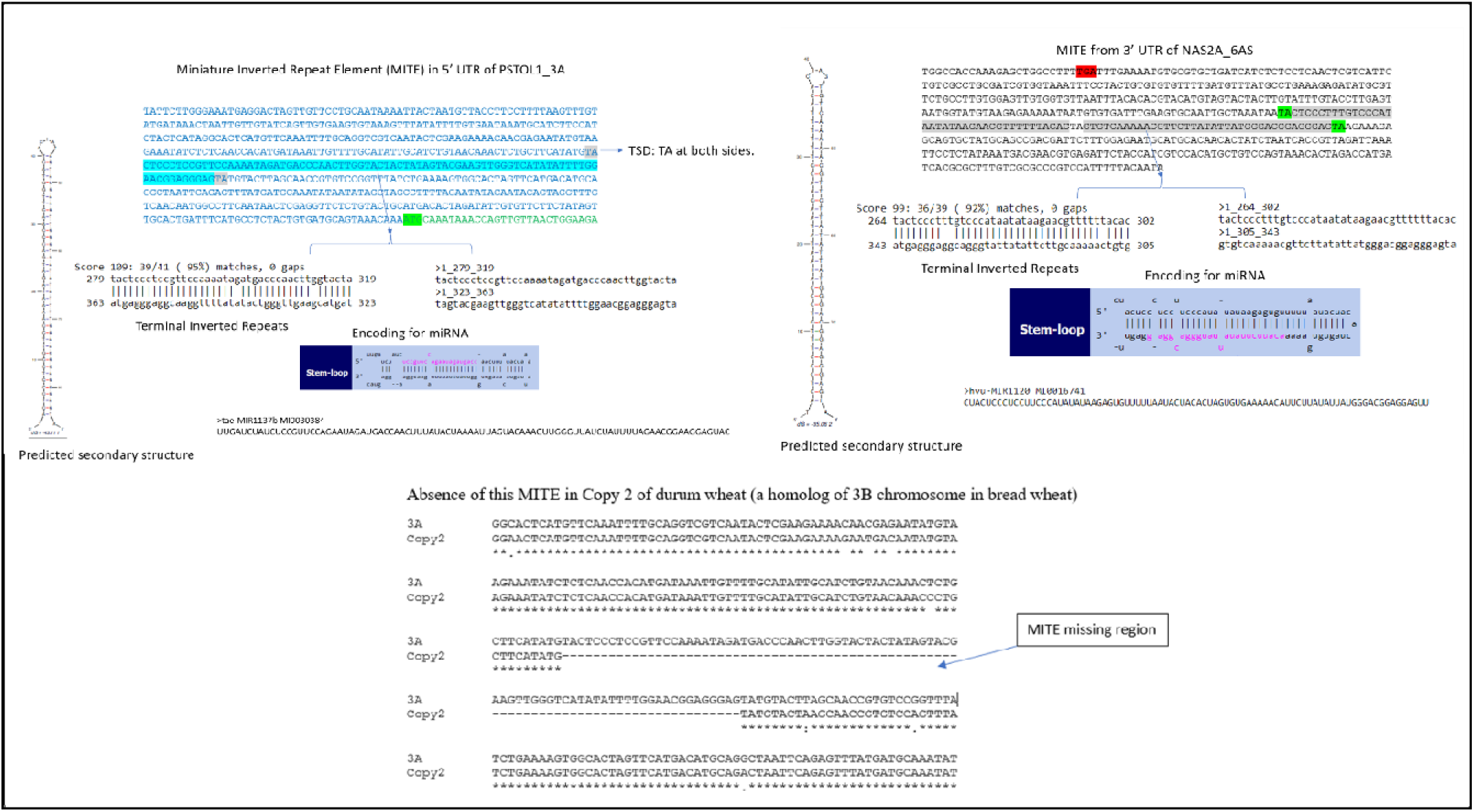
Prediction of MITES and MITES encoding for miRNA from specific homeologs and the comparison about the absence of MITE from a copy2 of durum wheat with a resemblance to 3B homeolog of bread wheat as the insertion only observed in 3A homeolog chromosome of bread wheat.

We identified 197 putative target genes for *tae-MIR1137b* including four protein kinases. However, there was no hit observed for either *TaPSTOL1_3DL* or *TaLrK10_3DL*, probably due to the absence of the MITE. This analysis suggests that this miRNA may have essential roles in the regulation of genes; however, it is unknown whether it regulates putative *TaPSTOL1-like* homeologs. Overall, our comparative analysis suggests that the third exon of *TaLrK10* and putative *TaPSTOL1-like* share high nucleotide sequence similarity in bread wheat. Further, the MITE insertion appears to have present in A and D subgenomes while absent in B subgenome of bread wheat.

### Phylogenetic and diversity analysis based on the protein sequences of putative TaPSTOL1-like and TaLrK10

The protein-based phylogenetic tree of putative *TaPSTOL1-like* and *TaLrK10* within *Poaceae* species indicate that all species studied with this phylogeny belonged to three clades (Fig. 2). The first clade consists of *OsPSTOL1* from *Oryzeae* species alone, suggesting the presence of *Oryzeae* species-specific haplotypes based on these two proteins (S4 Fig; e.g, “Y**K**G**E**LPNG**VPV**AVKMLEN” in *Oryzeae* species versus “Y**R**G**S**LPNG**REI**AVKMLKD” in *T.aestivum*_*Lr10K_* AAC49629_U51330). This clade clustered distantly to the second clade (see below). For instance, *OsPSTOL1* from *O.sativa* and *TaLrK10* from *T.aestivum* have 70% polymorphisms. Hence, *OsPSTOL1* from *Oryzae* species clustered distantly to those of *TaPSTOL1*/*TaLrK10* (clade II) of other tribes of *Poaceae* species.

The second clade comprises both *Triticeae* and other tribes of the *Poaceae* members due to a higher similarity between the aligned regions in the two proteins. For instance, “YRGDLSDGRQIAVKMLKD” from *TaPSTOL1*-3DL of bread wheat and “YRGGLSDGRQIAVKMLKD” from *AtLrK10* of *A.tauschii* differed by only 5.55% (S4 Fig). On the other hand, a lower distance between *TaPSTOL1*/*TaLrK10*, for instance, *LrK10* from *O.brachyantha* and *TaPSTOL1*-3DL from *T.aestivum* (19% variation) clustered together in clade II. In the clade III, only *TaLrK10* from *T.aestivum* appeared, which indicates that *TaPSTOL1* is distant to *TaLrK10* in bread wheat probably due to the absence of amino-terminal domain in *TaPSTOL1*. Further, this clade has also the *TaLrK10* sequence from chromosome 1AS (Genbank accession: Nucleotide_U51330, Protein_AAC49629). Therefore, the pattern of *TaLrK10* and *TaPSTOL1* gene divergences among the species studied appears to indicate two broad groups: while species in clade II have both *TaLrK10* (e.g. *RLKs*) and *TaPSTOL1* (e.g, *RLCKs*); the species in clade III appear to have limited to only *RLKs*. The overall pairwise mean distance through the JTT matrix-based model (Jones et al, 1992) suggests that both these proteins likely shared a common ancestral lineage. Overall, the polymorphic sites analysis revealed that *TaPSTOL1-like* has a higher nucleotide diversity compared to *TaLrK10* within *Triticeae*. However, it was unclear among the different tribe members of the *Poaceae* family (S3 Table).

The homogenous substitution through Monte Carlo 1000 replicates-based Disparity Index test (I_D_ Test) indicates that 12.38% of sites evolved with a pattern of homogenous substitutions with significant I_D_ Test. The estimated frequencies of amino acids for both genes ranged from 1.43% (W) to 9.11% (L). Substitution rate with 35 randomly selected sites (of 241 positions from 105 amino acid sequences), revealed that there are 13 conservative (C), ten semi-conservative (SC), and 12 non-conservative (NC) substitutions with the frequencies of 37.14%, 28.57%, and 34.29%, respectively. Sum of C and SC substitutions was 65.71%, which is higher than NC substitutions, suggesting that both genes share an evolutionary pattern of identity. These results indicate that both genes share a significant amount of sequence identity and hence are closely related to each other (S4 Table).

Protein-based divergence time tree indicated that the relative divergence time of *LrK10* from *Andropogoneae* and *Paniceae* species (Clade I) was earlier (0.57) (Fig. 4). Subsequently, *PSTOL1* and *LrK10* were diverged (0.47) in *Triticeae*, *Oryzeae*, *Brachypodieae*, *Andropogoneae* and *Paniceae* species (Clade II) with similar divergence times. The Tajima relative substitution rate analysis also supports the presence of both *PSTOL1* and *LrK10* in the clade II as these two genes share numerous monomorphic sites (250 sites) (S5 Table). Further analysis revealed a higher number of divergence sites and multiple sub-clusters comprising species involved in the divergence of both *PSTOL1*/*LrK10* from all four tribes. The *PSTOL1* most recently diverged in *Oryza* species (Clade III) (0.03). This analysis suggests that the environmental heterogeneity might have probably triggered a high divergence of *OsPSTOL1* in *Oryza species* [23]. This observation is also consistent with a lower pairwise nucleotide diversity noticed for *PSTOL1* (19%) compared to *LrK10* (30%). These results suggest that putative *PSTOL1* may be recently diverged compared to *LrK10* in the *Poaceae* tribe [9].

**Fig. 4.**
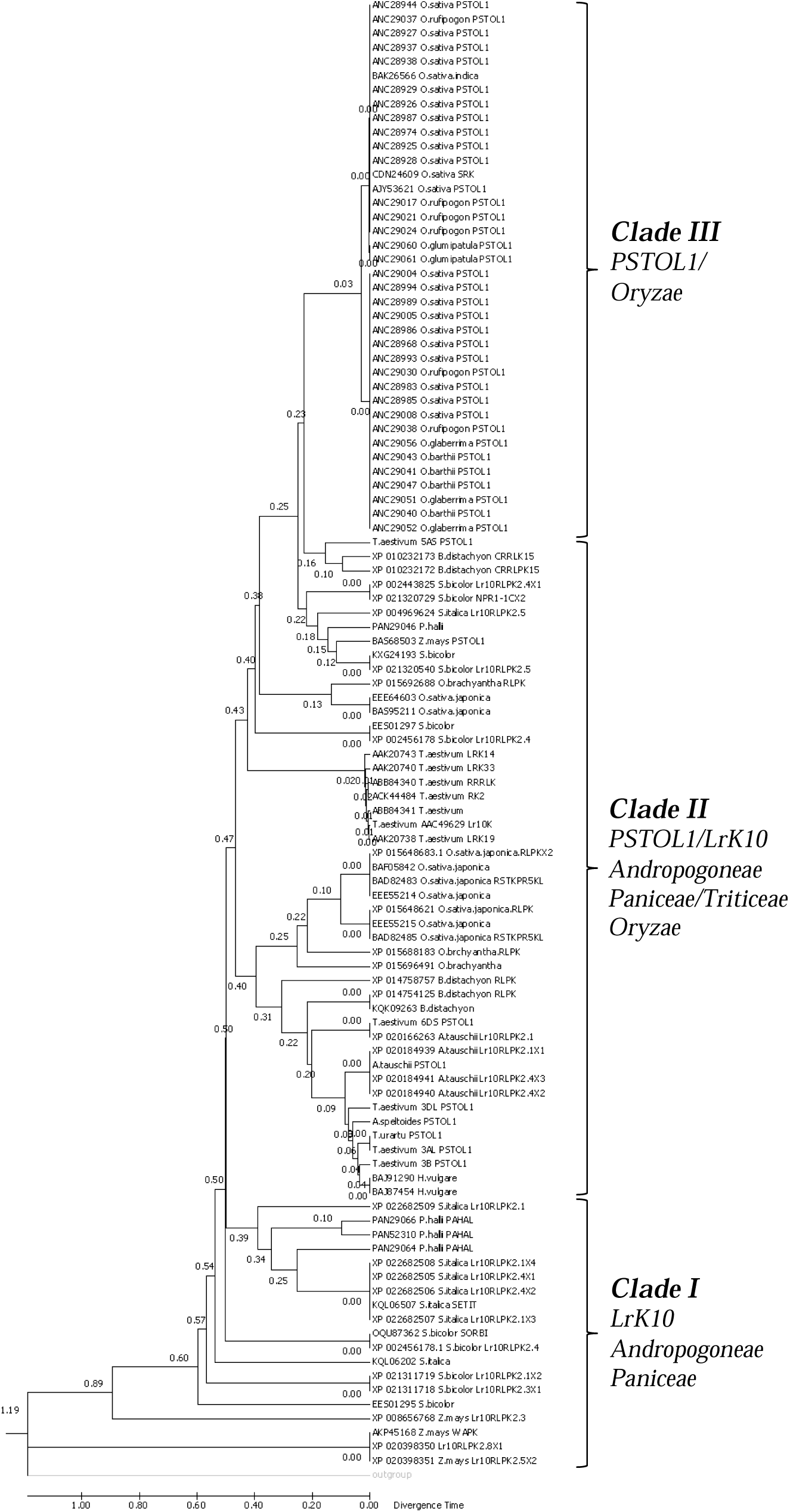
Relative Divergence time tree showing the relative divergence of *PSTOL1* and *LrK10* among diverse *Poaceae* species.

### Putative PSTOL1 related kinase search in different phyla

To understand the conservation of kinase related genes across different phyla, a search for putative *PSTOL1* kinase related protein sequences was done in SMART, and the Lifetree based representation is shown in Fig. 5. This analysis indicating the versatile conservation of this kinase related putative protein across different phyla represents the evolutionary importance of diversity and conservation of kinase related gene.

**Fig. 5.**
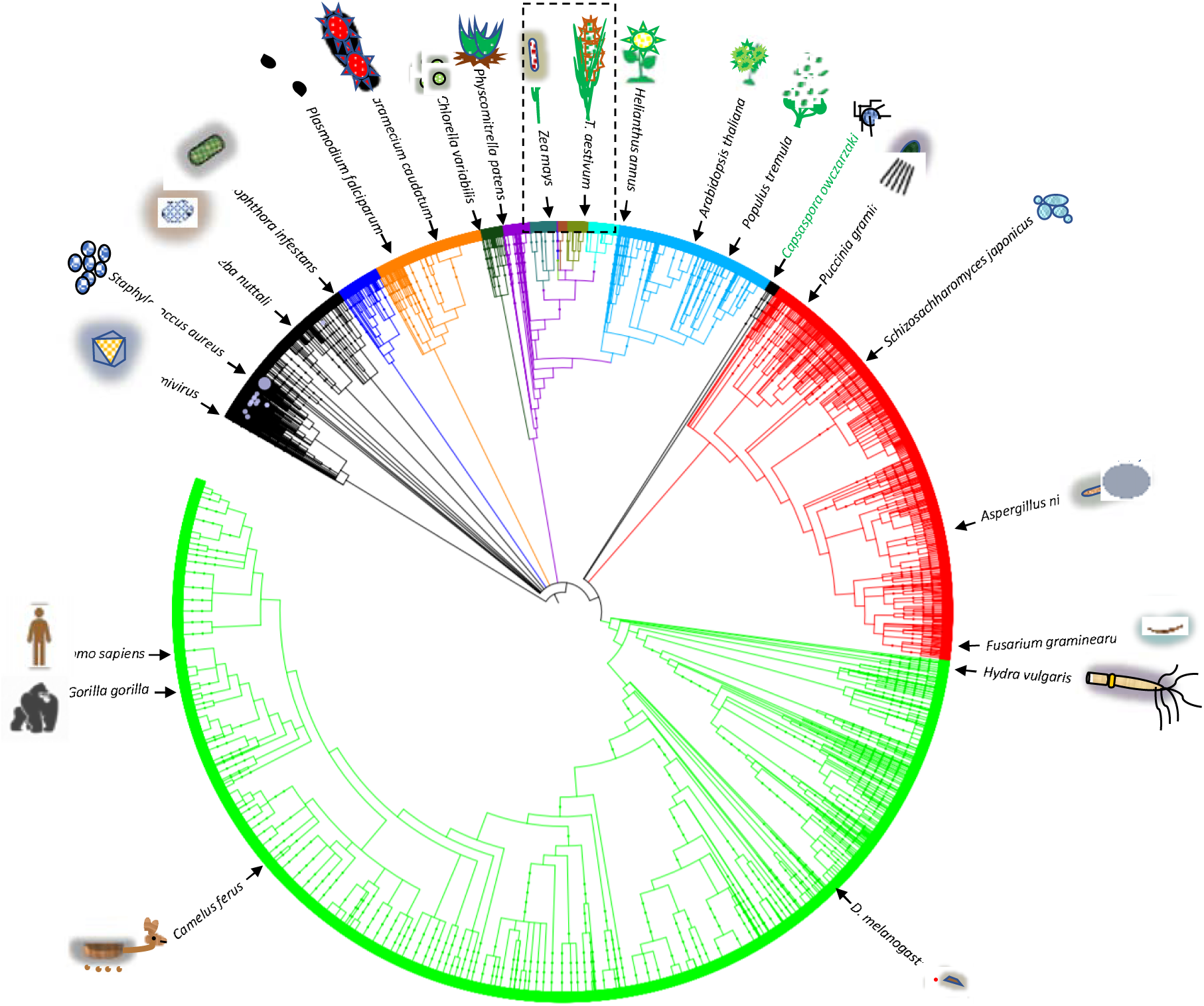
The life-tree based on putative *PSTOL1* related protein kinase sequences of across several phyla. The section of *Poaceae* is shown in the dotted-box. Orthologs of putative *PSTOL1* related protein kinase sequence from different phyla were predicted using the SMART (Simple Modular Architecture Research) tool and the life-tree was generated using iTOL.

### Comparison of putative TaPSTOL1 and TaLrK10 protein structures

We performed both motif and domain analyses for the protein coding region of the putative *TaPSTOL1* of all homeologs in bread wheat as well as the kinase aligned region of *AtLrK10* (kinase encoding part) from *A.tauschii*. The putative *TaPSTOL1*-3DL protein has 130 motifs, of which, 57 were exclusively kinase related, including three *CDK* (*Cyclin-Dependent Kinase*) motifs, one *MAPK* (*Mitogen-activated protein kinase*) motif, and one *PIKK* (*Phosphoinositide-3-OH-kinase-related kinases*) motif. The comparison between *PSTOL1* and *LrK10* among *Triticeae*, *Oryzeae*, *Andropogoneae,* and *Brachypodeae* tribes revealed several conserved motifs such as the CDK motif “RALIY” between positions 121-125, motif “RGLEY” between positions 162-166, except for the *LrK10* in bread wheat, and the motif “KMLL” between 336-339 positions. The putative *TaPSTOL1*-3DL from bread wheat showed a higher similarity with *AtLrK10* of *A.tauschii* in the kinase domain-encoding region (Data not shown). Genes belonging to the groups of *RLCKs*, including putative *TaPSTOL1*, do not possess the extracellular domain (Shiu and Bleecker, 2001) but do maintain a predicted transmembrane domain and kinase domain (S5 Fig).

We further studied the primary structural differences between putative *PSTOL1* and *LrK10* across several species of four tribes, which revealed the presence of 64 identical, 55 conservatively substituted and 12 semi-conservatively substituted residues. The kinase domain comprising 199 residues (starting at the position 52 in *TaPSTOL1*-3DL in bread wheat) revealed that there are 54 identical, 39 conservative and seven semi-conservative replacements when aligned with *TaLrK10*. The ATP binding signature “LGQGGFGAVYRGDLSDGRQIAVKM” starting at the position 59 was conserved between the two genes, with Gly17 conserved entirely across all the species studied. At the carboxyl (C) terminus, four identical and two conservative replacements were found between LrK10 and *PSTOL1* within the *Triticeae*, while the amino (N) terminal region was highly similar (S5 Fig).

The predicted ATP binding signature of Tyrosine kinase comprises 57 residues, starting from the position 86 and highly conserved between *PSTOL1* and *LrK10* with 12 identical residues, 17 conservatives and three semi-conservative changes. At the N-terminus, there were 23 residues entirely conserved between *AtLrK10* of *A.tauschii*, *B.distachyon* and *TaPSTOL1*-6DS of bread wheat. About 12 residues within the Ser/Thr active site signatures (starting at the position 175; “IVHFDIKPHNILL”) were highly conserved, with nine same sites and three conservative amino acid replacements between *PSTOL1* and *LrK10*. Further analysis using either *TaPSTOL1*-3DL or *TaLrK10* protein as a query in the Classification of Protein Domain Structures (CATH) server resulted in the identification of the Ser/Thr receptor kinase *PR5K* (E-value: 1.1e-75), which belongs to the group of phosphotransferase superfamily that catalyzes the transfer of a phosphate to the hydroxyl groups of their protein substrates. We further superimposed 148 representative domains within the phosphotransferase superfamily, which indicated that this domain is present across several species (Fig. S6).

The secondary structures of the putative *TaPSTOL1* and *TaLrK10* kinase domain regions are quite similar (Fig. 6). The putative *TaPSTOL1* vary from 51 - 67 α-helices while the kinase domain aligned region of *TaLrK10* had 39 α-helices between *T.aestivum* and *A.tauschii*, respectively. These regions also exhibited a higher frequency of identical amino acids, also consistently observed for extended beta-strands and random coils. The isoelectric focusing point (pI) varied from six to nine between the two genes when compared the homeologous sequences. The content of positively and negatively charged residues was similar between the two genes. The instability index estimated was 40 for both *TaPSTOL1* and the kinase domain-aligned region of *TaLrK10*. However, the instability index exceeded 40 when considered *TaLrK10* alone, which therefore appears to have a less stable protein compared to the putative *TaPSTOL1* (S6 Table).

**Fig. 6.**
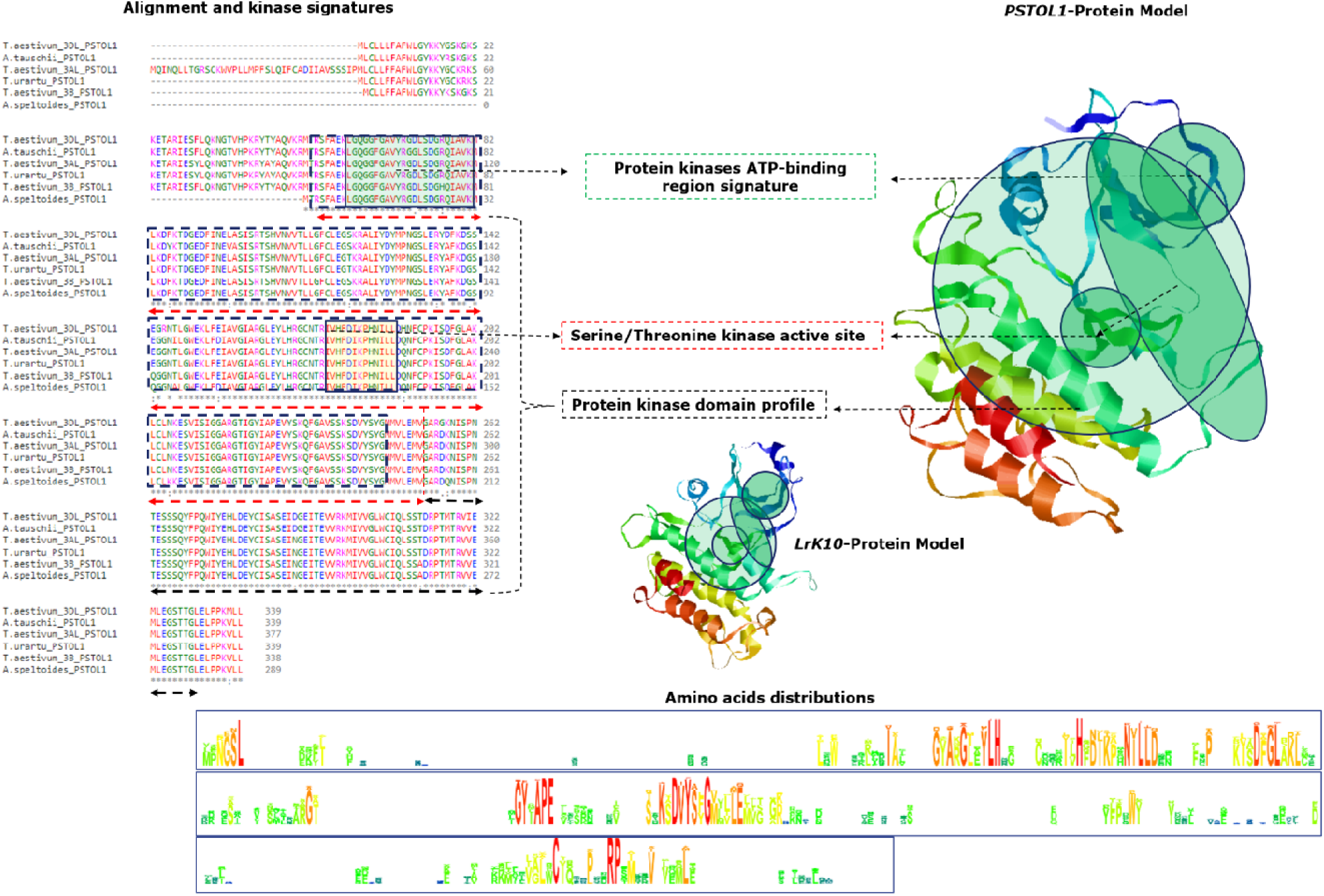
The protein 3D modelling of *PSTOL1* and *LrK10* and prediction of domains and aligned region of *PSTOL1* within bread wheat and its progenitors.

Protein-3D modelling revealed that the structural similarity in the kinase domain region between putative *TaPSTOL1* and *TaLrK10* is in accordance with the alignment of the two predicted proteins corresponding to their nucleotide sequence (Fig S6a). The *TaLrK10* model was created using the template *Brassinosteroid insensitive 1-associated receptor kinase 1* (PDB: 3ULZ.1), which showed 43.42% identity with the aligned kinase domain region.

The pattern of domains organization of putative *TaPSTOL1* homeologs within *Triticum* species (Fig. 7) suggests that it is similar to the *OsPSTOL1* of rice for not bearing the extracellular domain, which is typically found in *RLCKs* [19,59]. In contrast, the putative *LrK10* from *A.tauschii* and *T.aestivum* have this extracellular domain [60]. We further observed enormous structural rearrangements in the upstream region between putative paralogs *TaPSTOL1*_*5ASII* and *TaPSTOL1_5ASI* (*TaLrK10_5ASI*). A comparison between *LrK10* and *PSTOL1* homeologs in wheat reveals the presence of two patterns of structural organization with either single exon1 CDS (putative *PSTOL-like* gene) or three to four-exons with single or multiple transcripts with alternative splicing.

**Fig. 7.**
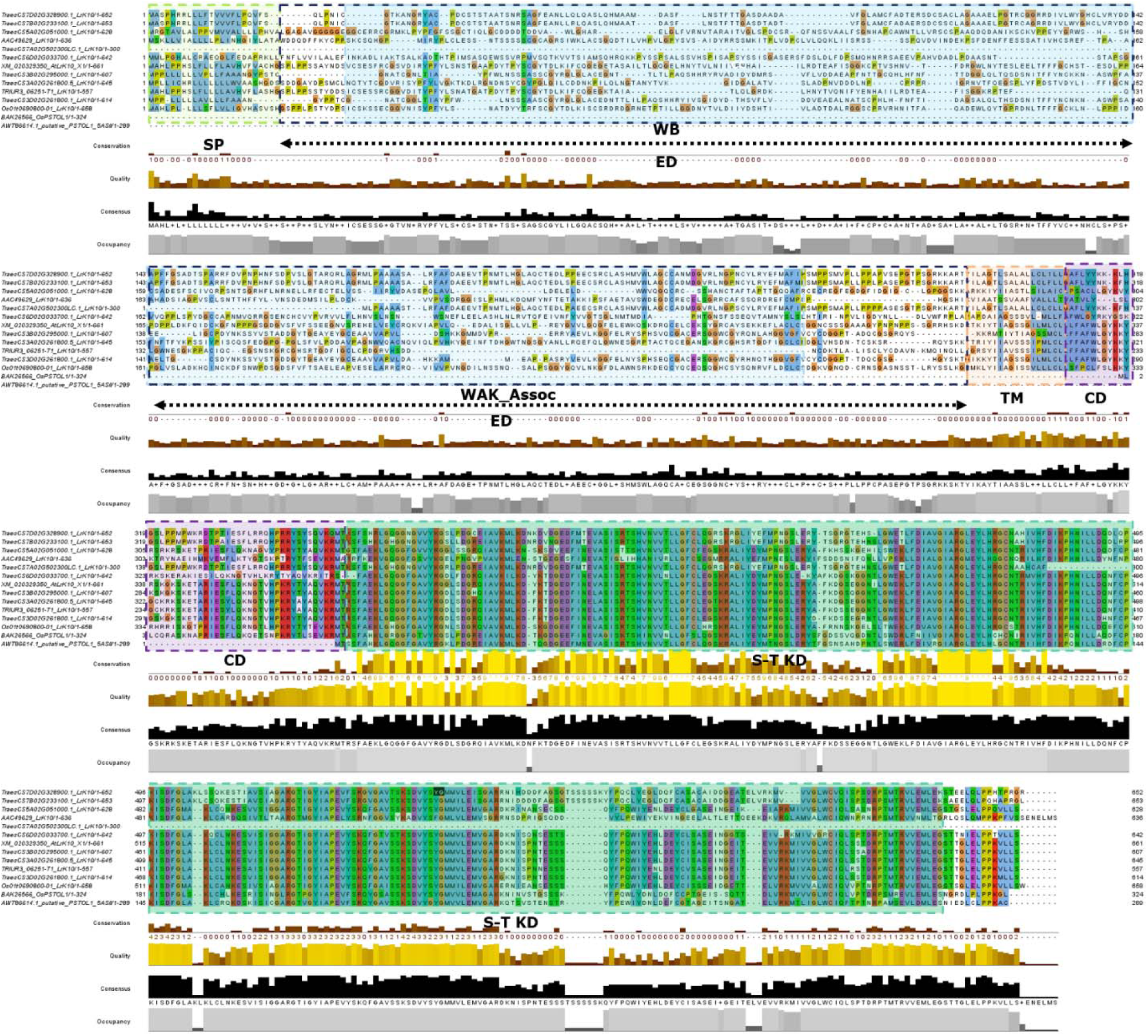
Protein domains structural organization and alignment between *LrK10* and *PSTOL1* gene orthologs and homeologs. SP: Signal Peptide, EB: Extra Cellular domain, TM: Transmembrane domain, CD: Conserved domain. S-T KD: Serine Threonine kinase domain. Domain predictions based on the source: Feuillet et al, 1997.

We noticed two distinct structural organization for *TaLrK10* in *Triticum* species (Fig. 8). In the wheat progenitor *A.tauschii*, *AtLrK10* (Genbank accession: XP_020184939) carry a partial *wall-associated protein kinase* carboxyl region (*WAK*; 94 amino acids) in the N-terminus, which is comparable across diverse *Poaceae* species including sorghum [14]. However, *WAK* domain appears to have absent in some of the *TaLrK10* homeologs in bread wheat (chromosomes 6 and 7) including *TaLr10* (Genbank accession: U51330) [18]. To understand better with the organization of domains in putative *PSTOL1* gene homeologs and orthologs, we have further depicted an illustration based on the prediction model of *LrK10* [18]. The alignment-based prediction model revealed the higher conservation of Serine-Threonine Kinase domain in *PSTOL1* and its related *LrK10* protein.

**Fig. 8.**
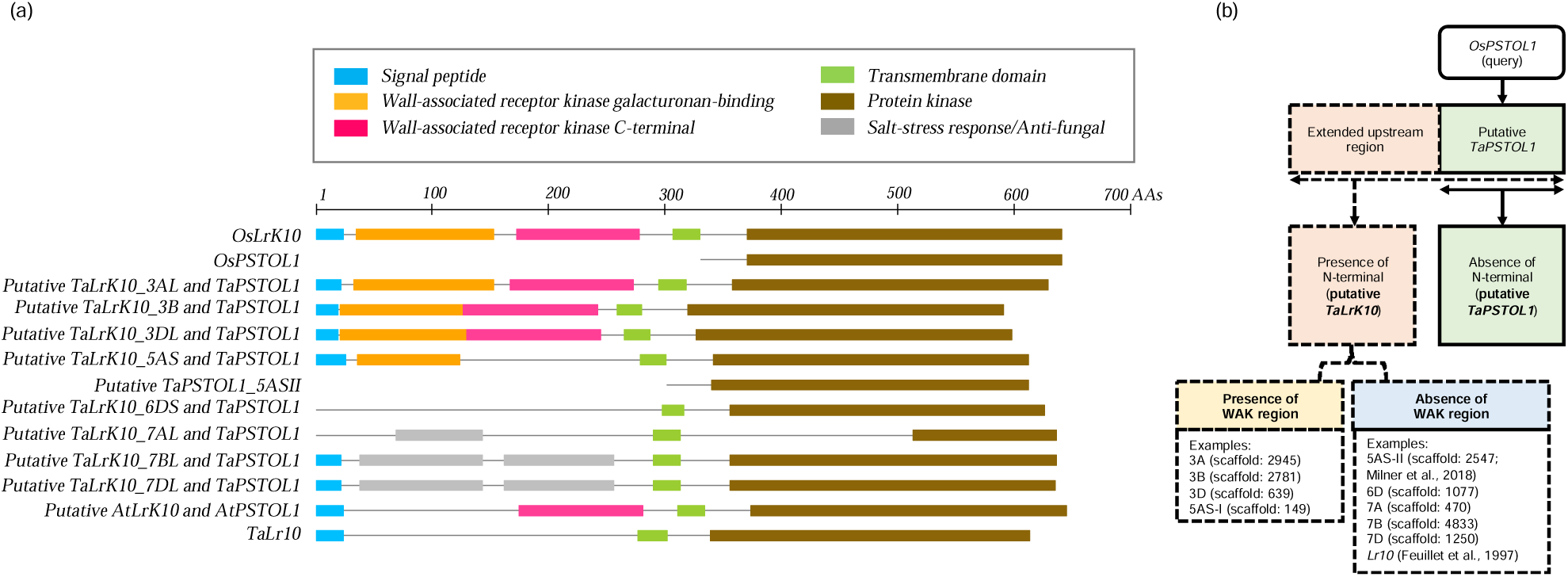
Structural organization of putative *TaLrK10* and *TaPSTOL1* genes in *Triticum* species. (a) The organization, presence, and absence of different domains in putative *TaLrK10* and *TaPSTOL1* in *Triticum* species. The amino-terminal domain is present in putative *TaLrK10* but absent in *TaPSTOL1*. Similarly, the *WAK* (wall-associated kinase) is observed in putative *LrK10* in *A.tauschii*, and few homeologous putative *TaPSTOL1* in bread wheat while absent in all other homeologs. (b) Schematic representation of putative *TaLrK10* and *TaPSTOL1-like* genes, and the presence or absence of N-terminal domain, and *WAK* region in wheat.

## Discussion

### Commonalities and dissimilarities of putative TaPSTOL1-like and TaLrK10 gene sequences

Both *OsPSTOL1* and *TaLrK10* genes have been known to play different functional roles in plants. However, it is unknown whether they belong to the same ancestral lineage. Using a comparative bioinformatic framework, we examine that both the putative *TaPSTOL1-like* and *TaLrK10* genes are closely related to each other, likely derived from a common ancestral origin. Based on BLASTP and BLASTN, eight putative homeologs of *TaPSTOL1-like* were detected in bread wheat, consistent with several *PSTOL1* homologs/orthologs identified in other species such as sorghum (14) and maize [15]. Similar to *OsPSTOL1*, all eight putative wheat *TaPSTOL1*-like proteins are predicted to have a conserved Ser/Thr kinase domain. In contrast to our study, a recent study reported a single copy of *TaPSTOL1* on chromosome 5A in wheat [16]. Such a difference in the number of homeologs identified between these two studies could potentially be related to the incomplete (as well as continuous update) nature of the wheat genome database and germplasm used.

While the putative *TaPSTOL1-like* homeologs identified based on the BLAST in our study were not functionally tested, we further leveraged wheat expression database [56] to show that all the putative *TaPSTOL1-like* homeologs identified in our study showed enhanced expression under low-P as compared to control conditions, suggesting that all these putative homeologs could be expressed under low-P. Consistently, the expression of the *TaPSTOL1* like gene was observed (on chromosome 5AS; Acc# MH043199) in root tips and root hairs under different P concentrations [16]. The putative *TaPSTOL1-like* homeologs identified in this study could therefore be analogous to previously reported *PSTOL1* genes in maize [14,15] and rice [9,11,12,13,23].

It is interesting to note that the conserved kinase region appears to have a high nucleotide similarity (96%) with the third exonic region of *TaLrK10_3DL* from bread wheat. The multiple-sequence alignment (*TaLrK10_3DL* and *TaPSTOL1-3DL*) further indicated that the putative *TaPSTOL1_3DL* overlaps with the genomic locations of *TaLrK10_3DL*. Indeed, the putative *TaPSTOL1* appears to have derived from the last exonic region (3^rd^ exon) of *TaLrK10_3DL* in bread wheat that indeed showed more than 70% of deletions relative to its ancestral *AtLrK10* from *A.tauschii*. While the putative *TaPSTOL1_3DL* appears to have lacking the other exons that encode the extracellular domain in *TaLrK10_3DL*, as was also observed in rice [9], these comparative analyses inform that third exon of *TaLrK10_3DL* (3DL: 363607922 bp to 363608939 bp) would indeed act as a putative single exon-based *TaPSTOL1-like* gene in bread wheat. Therefore, the putative *TaPSTOL1* (*RLCK*) seems to have the variant of typical *TaLrK10* in bread wheat that belong to *RLK* the family [61].

### Both PSTOL1 and LrK10 have conserved kinase domains

The protein kinase family is a large dynamic group with highly conserved functional domains and motifs [62,63]. Hence, we focused our comparative analysis on the relevancy of the different structural organization of domains and motifs. For instance, the motif “RAILY” is highly conserved between *PSTOL1* and *LrK10* across diverse species within the *Poaceae*. There were, however, some subtle differences detected at either the N-terminal or the C-terminal level and also observed for the RGLEY and KMLL motifs. The kinase domain was highly conserved with many identical amino acids, and the ATP binding signature had 23 out of 24 amino acids highly conserved, indicating a robust structural redundancy in both the analyzed proteins.

### Structural analysis of TaLrK10 indicates two distinct patterns in Triticum species

The predicted physicochemical properties indicate the secondary and tertiary structures of the kinase domains of putative *TaPSTOL1-like* and *TaLrK10* are similar in the theoretical isoelectric point and instability index. It has previously been shown an *in-vitro* phosphorylation activity for *OsPSTOL1* using recombinant technology [9]. Consistent with this study, our computational analysis also indicates that *TaPSTOL1* and *TaLrK10* are closely related to the superfamily transferase (*Phosphotransferase*) domain 1, and is also closely associated with nonspecific *serine-threonine protein kinases* (2.7.11.1) and *serine-threonine protein kinases* (2.7.10.1), which are part of the CATH superfamily cluster number 12. Our analysis, therefore, indicates that the putative *TaPSTOL1-like* is structurally more similar to *AtLrK10-L2* of *A.tauschii* than to *TaLrK10* of *T.aestivum*.

We observed two distinct structural organization for putative *TaLrK10* within *Triticum* species. *TaLrK10* contains a wall-associated kinase (*WAK)* carboxyl region in at C-terminus in the wheat progenitor *A.tauschii*, as was also predicted in some *SbPSTOL* homologs in sorghum [14]. However, such a *WAK* region is absent in the majority of the putative *TaLrK10* homeologs predicted in bread wheat. The *WAK* proteins possess a typical cytoplasmic Ser/Thr kinase signature, which typically carry an extracellular domain (e.g. *LrK10* protein), mediating cross-talk with environmental signals [14,64,65,]. It is known that chitin from fungal pathogens [66,67] and pathogen-based induction of salicylic acid trigger *WAK* containing genes for the survival of host plants [68]. The apparent absence of such a *WAK* region in majority of the putative *TaLrK10* homeologs (particularly *TaLr10* from 1A; [18] in bread wheat supports the prevailing notion that the *TaLrK10* gene is no longer considered to the ortholog of *Lr10* gene in bread wheat although it may still encode a kinase domain involved in plant-pathogen interactions [69]. We suspect that *TaLrK10* lacking the *WAK* region might have served as the basis for the divergence of the putative *TaPSTOL1-like* gene as a single-exon based gene without the *WAK* region while retaining the kinase domain. The putative *TaPSTOL1-like*, which lacks the amino-terminal domain and thus is structurally similar to the third exon of *TaLrK10* in bread wheat. Further research is needed to validate the presence or absence of the *WAK* region likely associated with different functional roles of RLK proteins in plants.

### Conclusion

Within the comparative genomics framework, our study examines that both the putative *TaPSTOL1* and *TaLrK10* genes are evolutionarily related to each other. However, the divergence of *TaLrK10* occurred earlier than the putative *TaPSTOL1* evolved. Each gene appears to have slightly modified their functional properties such as N-terminus and the extracellular domain, and there is an insertion of new MITE in 5′UTR of the putative *TaPSTOL1*. The sharable properties of their amino acids indicate that most of the mutations between these two genes could be synonymous and gene conversion supports the presence of tracts between these two genes among *Poaceae* species. Studying the evolutionary and functional aspects of the putative *TaPSTOL1* and *TaLrK10* through gene-knockout models would further facilitate the comprehensive understanding of their functional roles in plants.

## Acknowledgement

Authors duly acknowledge the support provided by the International Wheat Yield Partnership (IWYP) through the project number IWYP39 and the United States Agency for International Development (W0293.01.01 to RV at CIMMYT, Mexico), and the first author, KT, carried out the work under the guidance of RV during his tenure at CIMMYT, Mexico. Corresponding authors are grateful to Yogendra Kalenahalli, Stuart Roy, and Sigrid Heuer for their valuable suggestions.

## Author contributions

Plan the work, analyzed the data and wrote the manuscript: KT and RV. Assisted in revision of manuscript: CP, CK, DS, GT, DS, DT, AT, PV.

## Declaration

The authors have declared no competing interest.

## Supplementary Information

**S1 Fig.**
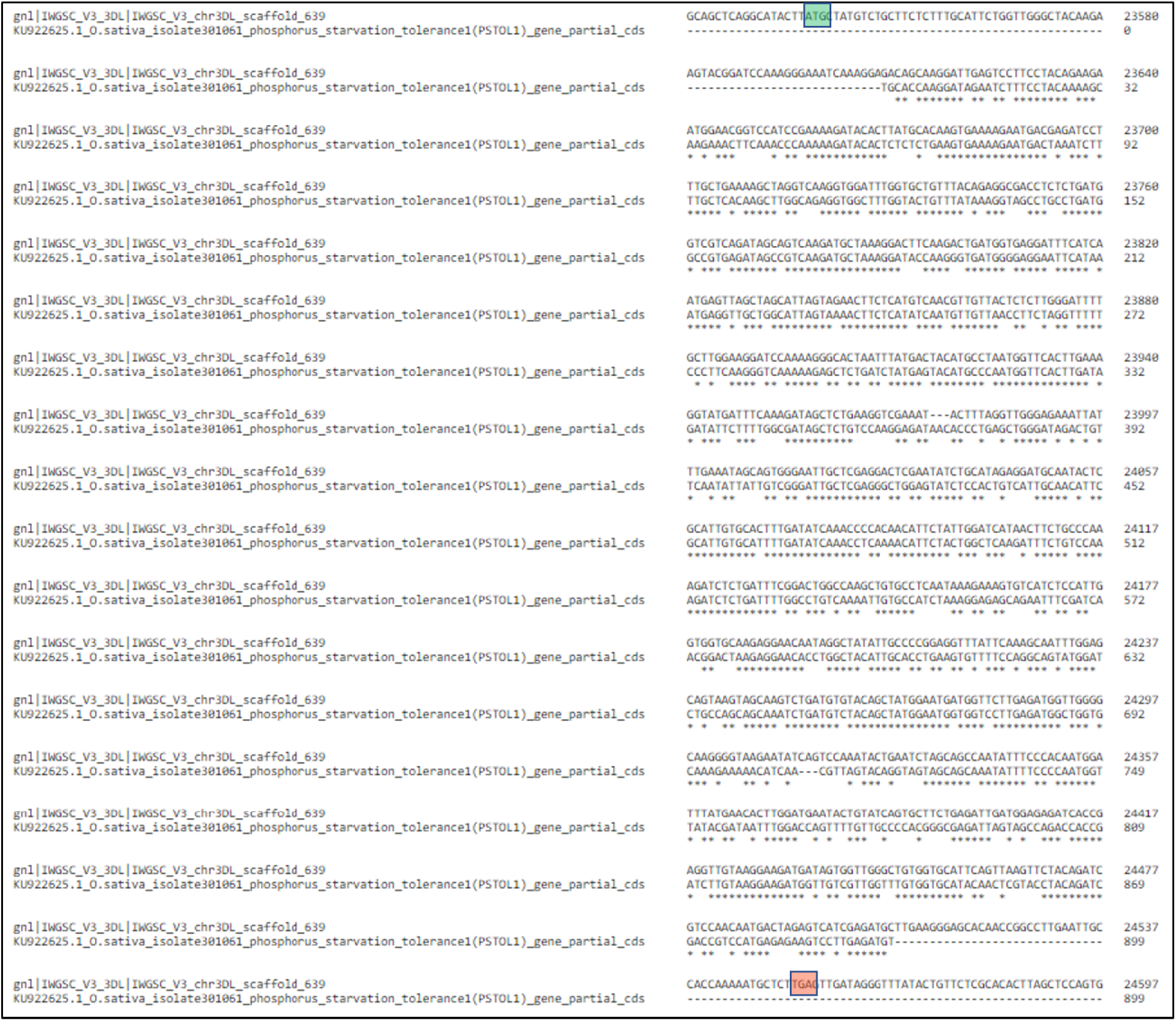
Alignment between *O. sativa PSTOL1* gene and *PSTOL1* gene from 3DL chromosome of bread wheat.

**S2a Fig.**
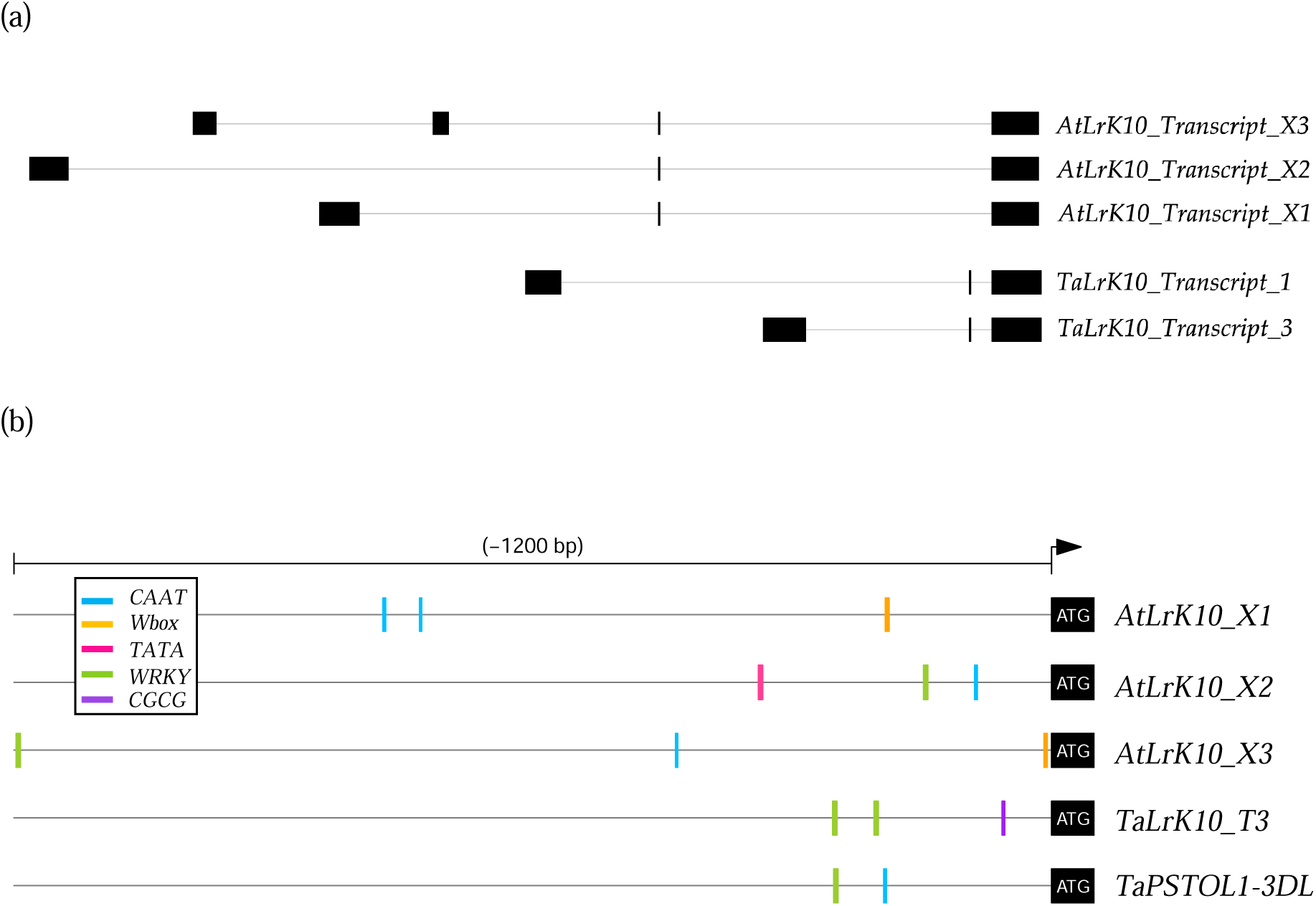
Gene comparison between *LrK10* and *PSTOL1* in bread wheat and it’s wild relative *Aegilops tauschii*. Different putative transcript variants of *LrK10* in *A.tauschii*, bread wheat and putative *PSTOL1-like* in bread wheat. Transcript variant X3 in *A.tauschii* has four predicted exons while all other transcripts in *A.tauschii* and bread wheat have three predicted exons. The conserved kinase domain across these multiple transcripts can be noticeable (the last exon in all transcripts). The black-filled rectangles indicate exons while horizontal grey-colored lines represent introns. **S3b Fig.** Predicted promoter transcription factor binding motifs in the upstream region (−1200 bp) for transcripts of *AtLrK10* and *TaLrK10* (Chromosome 3D).

**S3 Fig.**
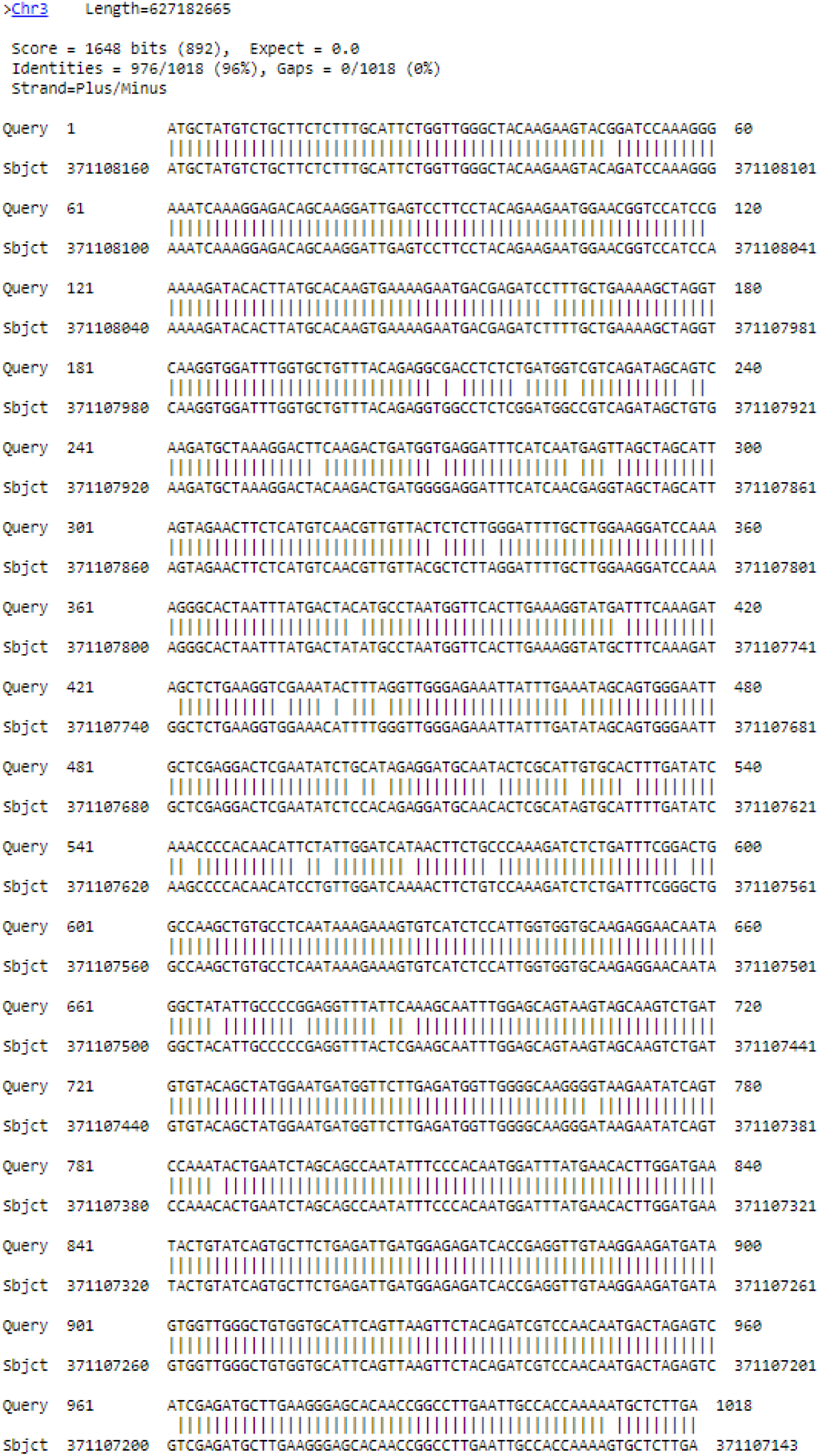
*PSTOL1* alignment from chromosome group 3 of *A. tauschii* and *PSTOL1*-3DL from bread wheat.

**S4 Fig.**
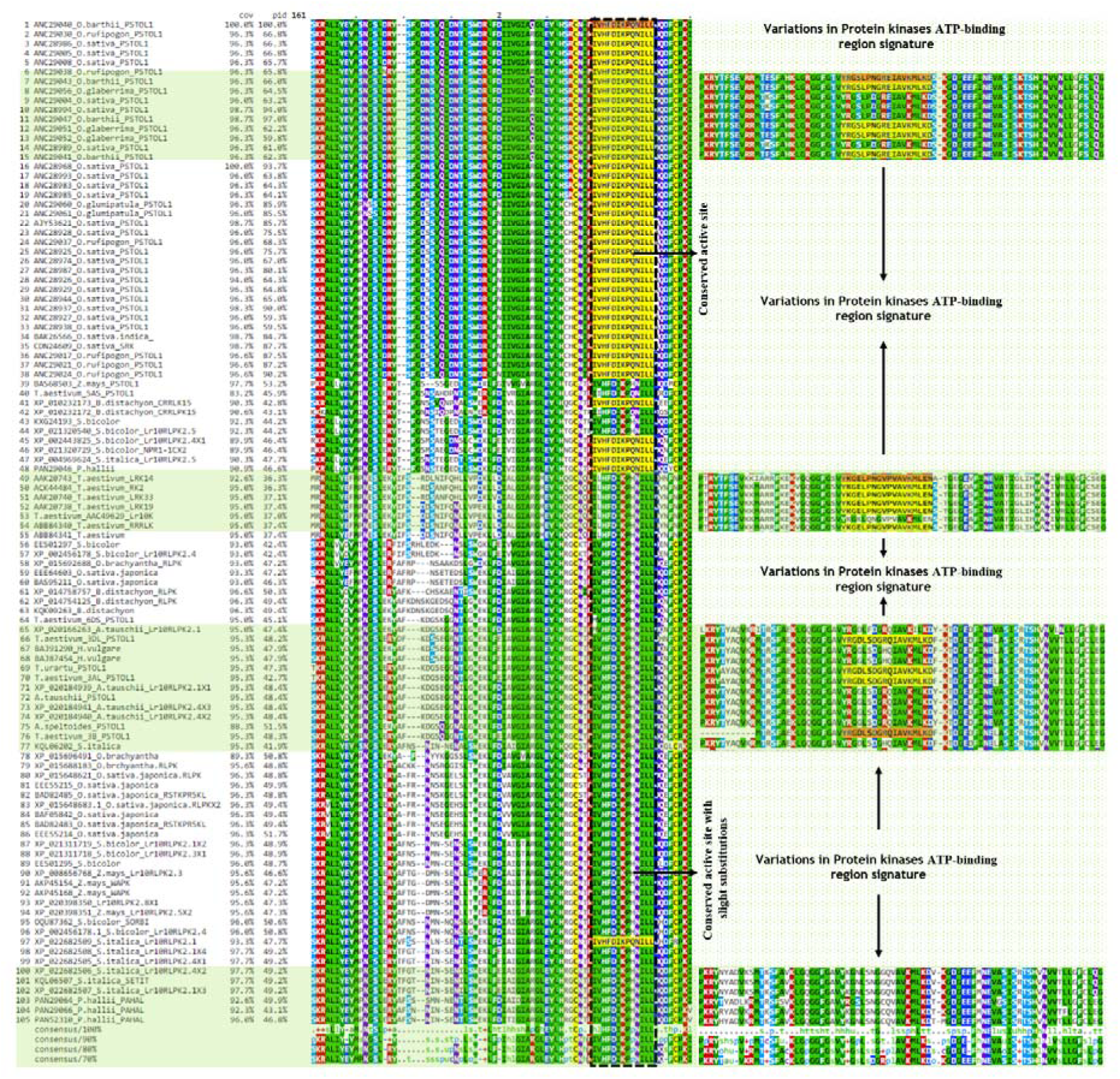
Alignment showing similarities among the conserved region alignment from different species of *poaceae* between *PSTOL1* gene and *LrK10* gene members.

**S5 Fig.**
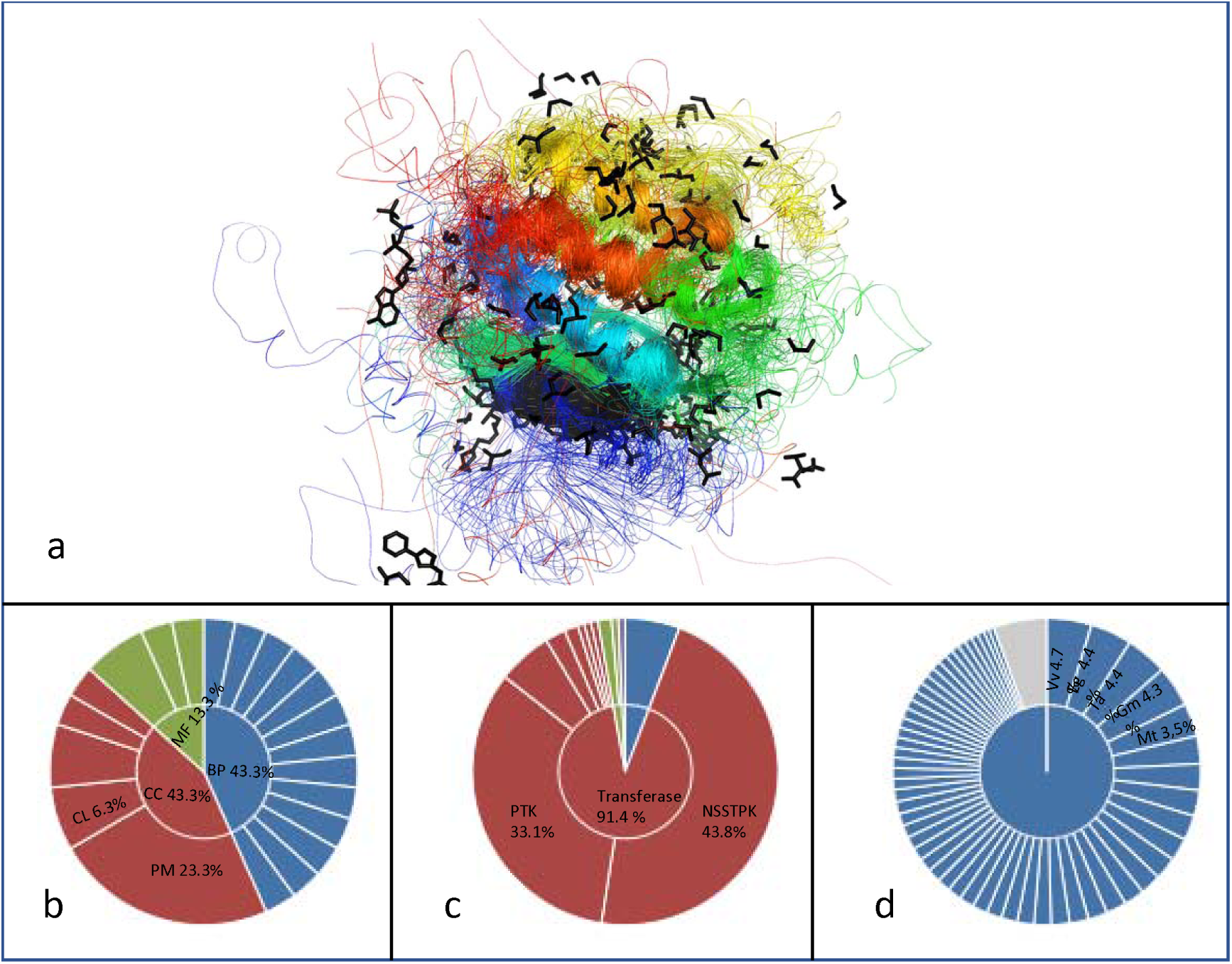
Superposition of 148 representative kinase domains within the superfamily Transferase (Phosphotransferase) domain 1 (a); Gene ontology indicating the percentage of functional properties of this superfamily (b); Enzyme commission numbers showing the percentage of non-specific serine/threonine protein kinase within this superfamily (c); and observation of the percentage of species diversity concerning this superfamily (d). BP: Biological Process, CL: Cytosol, CC: Cellular Components, MF: Molecular Function.

**S6 Fig.**
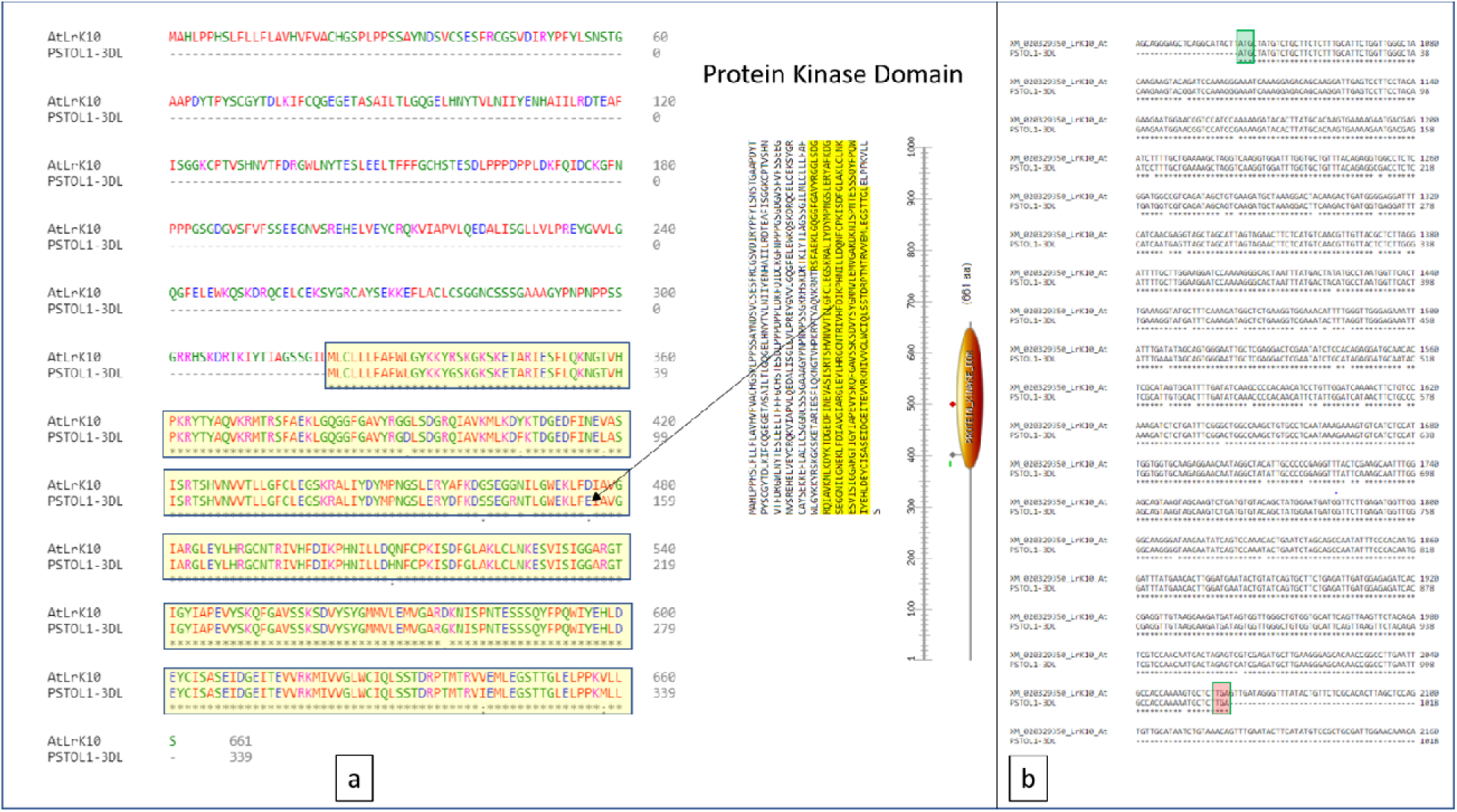
(a) The protein alignment of *PSTOL1*-3DL from bread wheat and *LrK10* gene from *A. tauschii* on their kinase domain. (b) Aligned regions of nucleotide sequences covering region of *PSTOL1* gene with *LrK10* gene.

**S1 Table.**
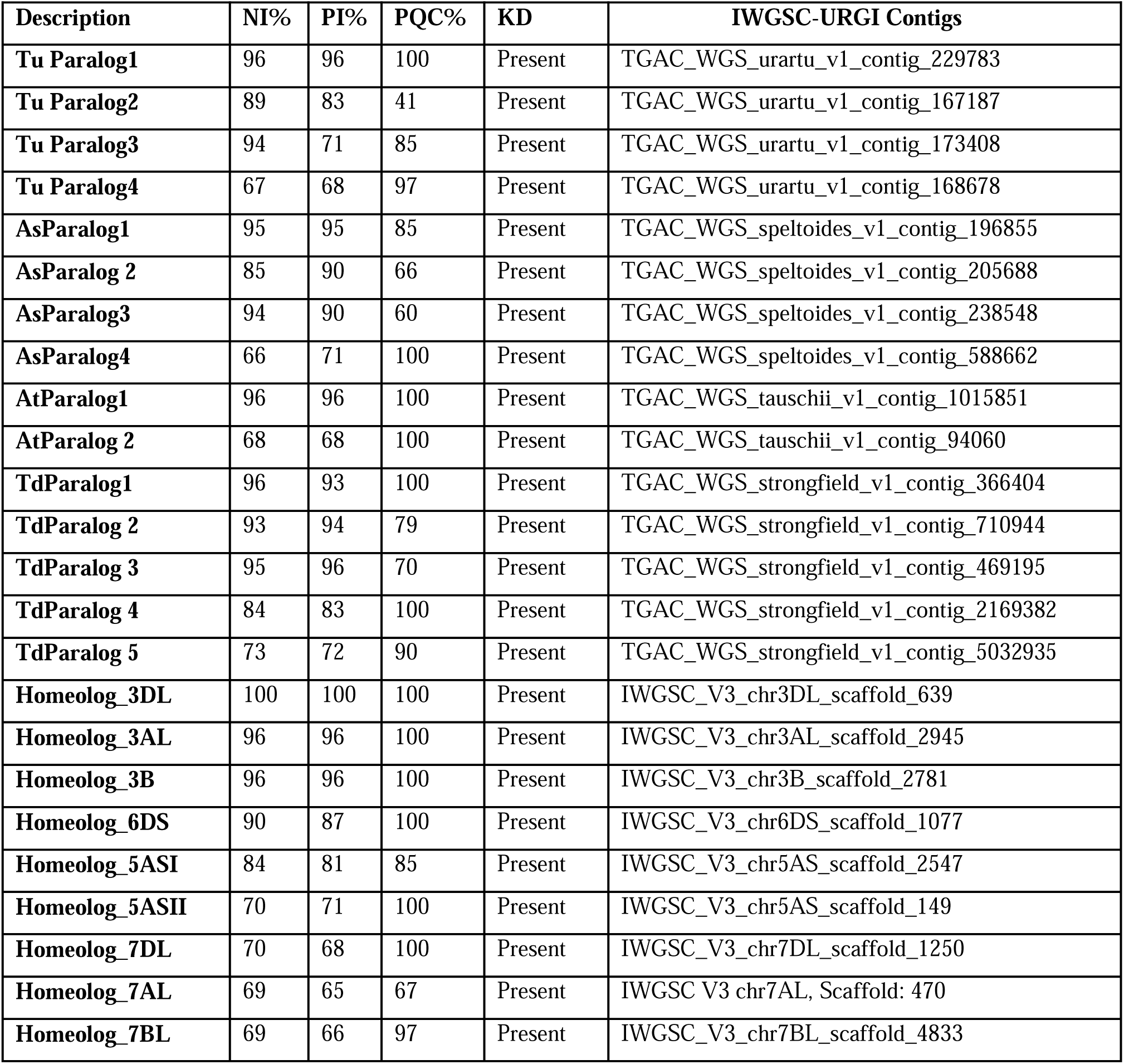
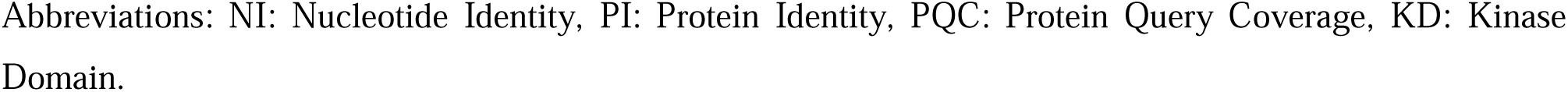
Identified orthologs/paralogs of *PSTOL1* gene in wheat wild progenitors and homeologs of durum wheat and bread wheat and contig IDs.

| Description | NI% | PI% | PQC% | KD | IWGSC-URGI Contigs |
| --- | --- | --- | --- | --- | --- |
| **Tu Paralog1** | 96 | 96 | 100 | Present | TGAC_WGS_urartu_v1_contig_229783 |
| **Tu Paralog2** | 89 | 83 | 41 | Present | TGAC_WGS_urartu_v1_contig_167187 |
| **Tu Paralog3** | 94 | 71 | 85 | Present | TGAC_WGS_urartu_v1_contig_173408 |
| **Tu Paralog4** | 67 | 68 | 97 | Present | TGAC_WGS_urartu_v1_contig_168678 |
| **AsParalog1** | 95 | 95 | 85 | Present | TGAC_WGS_speltoides_v1_contig_196855 |
| **AsParalog 2** | 85 | 90 | 66 | Present | TGAC_WGS_speltoides_v1_contig_205688 |
| **AsParalog3** | 94 | 90 | 60 | Present | TGAC_WGS_speltoides_v1_contig_238548 |
| **AsParalog4** | 66 | 71 | 100 | Present | TGAC_WGS_speltoides_v1_contig_588662 |
| **AtParalog1** | 96 | 96 | 100 | Present | TGAC_WGS_tauschii_v1_contig_1015851 |
| **AtParalog 2** | 68 | 68 | 100 | Present | TGAC_WGS_tauschii_v1_contig_94060 |
| **TdParalog1** | 96 | 93 | 100 | Present | TGAC_WGS_strongfield_v1_contig_366404 |
| **TdParalog 2** | 93 | 94 | 79 | Present | TGAC_WGS_strongfield_v1_contig_710944 |
| **TdParalog 3** | 95 | 96 | 70 | Present | TGAC_WGS_strongfield_v1_contig_469195 |
| **TdParalog 4** | 84 | 83 | 100 | Present | TGAC_WGS_strongfield_v1_contig_2169382 |
| **TdParalog 5** | 73 | 72 | 90 | Present | TGAC_WGS_strongfield_v1_contig_5032935 |
| **Homeolog_3DL** | 100 | 100 | 100 | Present | IWGSC_V3_chr3DL_scaffold_639 |
| **Homeolog_3AL** | 96 | 96 | 100 | Present | IWGSC_V3_chr3AL_scaffold_2945 |
| **Homeolog_3B** | 96 | 96 | 100 | Present | IWGSC_V3_chr3B_scaffold_2781 |
| **Homeolog_6DS** | 90 | 87 | 100 | Present | IWGSC_V3_chr6DS_scaffold_1077 |
| **Homeolog_5ASI** | 84 | 81 | 85 | Present | IWGSC_V3_chr5AS_scaffold_2547 |
| **Homeolog_5ASII** | 70 | 71 | 100 | Present | IWGSC_V3_chr5AS_scaffold_149 |
| **Homeolog_7DL** | 70 | 68 | 100 | Present | IWGSC_V3_chr7DL_scaffold_1250 |
| **Homeolog_7AL** | 69 | 65 | 67 | Present | IWGSC_V3_chr7AL, Scaffold: 470 |
| **Homeolog_7BL** | 69 | 66 | 97 | Present | IWGSC_V3_chr7BL_scaffold_4833 |

Abbreviations: NI: Nucleotide Identity, PI: Protein Identity, PQC: Protein Query Coverage, KD: Kinase Domain.

**S2 Table.** MITE (Miniature Inverted Repeat Transposable Element) from 5’ UTR of wheat wild progenitors, durum wheat and bread wheat.

| **Species** | Presence or absence<br>of the *PSTOL1* gene | Presence or absence of the MITE at 5' UTR of this<br>*PSTOL1* gene |
| --- | --- | --- |
| *T. urartu* | Present | Present |
| *A. speltoides* | Present | Absent |
| *A. tauschii* | Present | Absent |
| *T. monococcum* | Present | Absent |
| *A. sharonensis* | Present | Absent |
| *T. durum* |  |  |
| Copy1_3AL ortholog | Present | Present |
| Copy2_3B Ortholog | Present | Absent |
| *T. aestivum* |  |  |
| Copy1_ Chromosome 3AL | Present | Present |
| Copy2_ Chromosome 3B | Present | Absent |
| Copy3_ Chromosome 3DL | Present | Present with many mutations |

**S3 Table.**
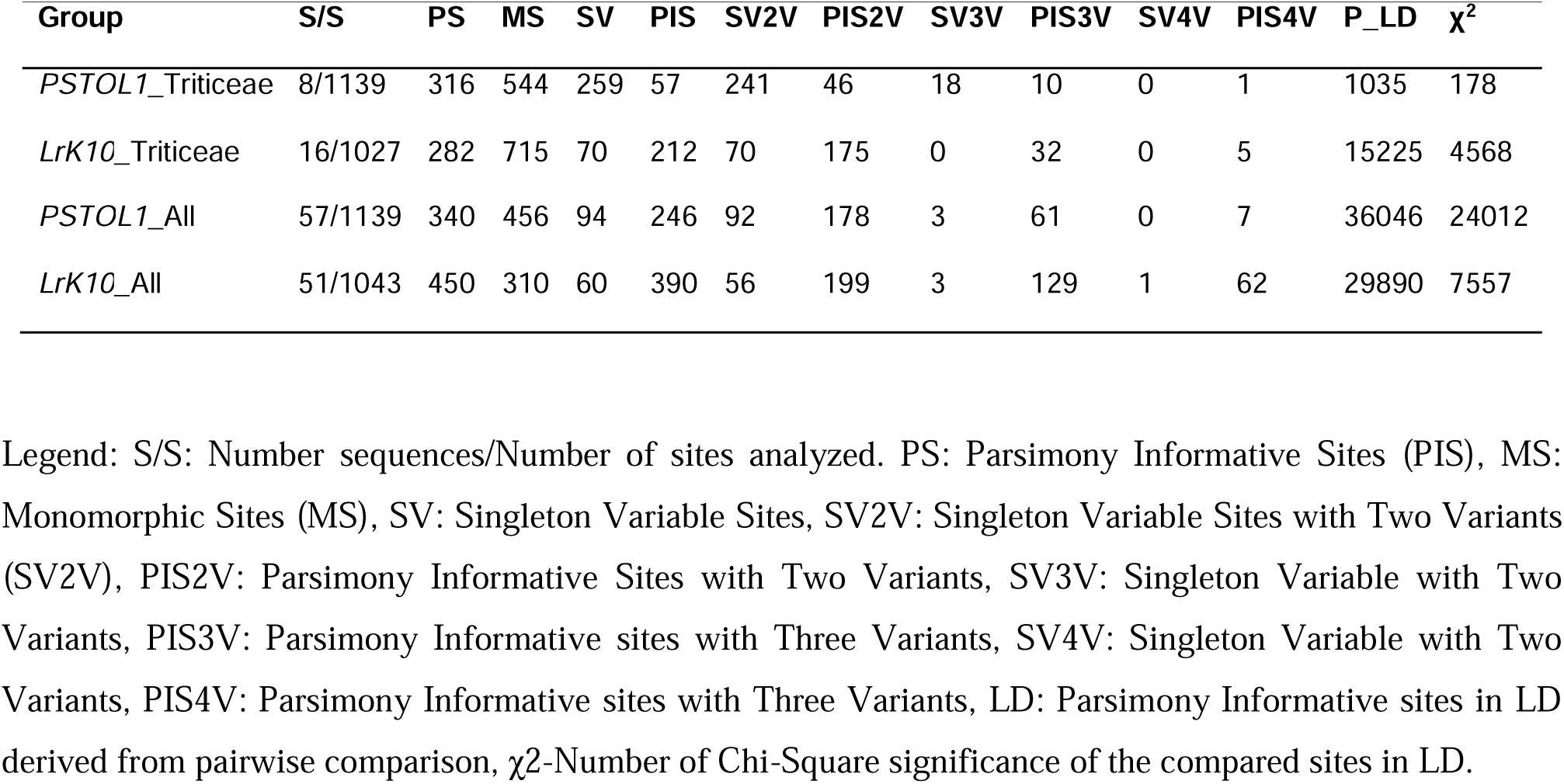
Parsimony Informative Sites (PIS) and other sites in each group of the genes from diverse species.

| Group | S/S | PS | MS | SV | PIS | SV2V | PIS2V | SV3V | PIS3V | SV4V | PIS4V | P_LD | $\chi^2$ |
| --- | --- | --- | --- | --- | --- | --- | --- | --- | --- | --- | --- | --- | --- |
| *PSTOL1*_Triticeae | 8/1139 | 316 | 544 | 259 | 57 | 241 | 46 | 18 | 10 | 0 | 1 | 1035 | 178 |
| *LrK10*_Triticeae | 16/1027 | 282 | 715 | 70 | 212 | 70 | 175 | 0 | 32 | 0 | 5 | 15225 | 4568 |
| *PSTOL1*_All | 57/1139 | 340 | 456 | 94 | 246 | 92 | 178 | 3 | 61 | 0 | 7 | 36046 | 24012 |
| *LrK10*_All | 51/1043 | 450 | 310 | 60 | 390 | 56 | 199 | 3 | 129 | 1 | 62 | 29890 | 7557 |

Legend: S/S: Number sequences/Number of sites analyzed. PS: Parsimony Informative Sites (PIS), MS: Monomorphic Sites (MS), SV: Singleton Variable Sites, SV2V: Singleton Variable Sites with Two Variants (SV2V), PIS2V: Parsimony Informative Sites with Two Variants, SV3V: Singleton Variable with Two Variants, PIS3V: Parsimony Informative sites with Three Variants, SV4V: Singleton Variable with Two Variants, PIS4V: Parsimony Informative sites with Three Variants, LD: Parsimony Informative sites in LD derived from pairwise comparison, $\chi^2$ -Number of Chi-Square significance of the compared sites in LD.

**S4 Table.**
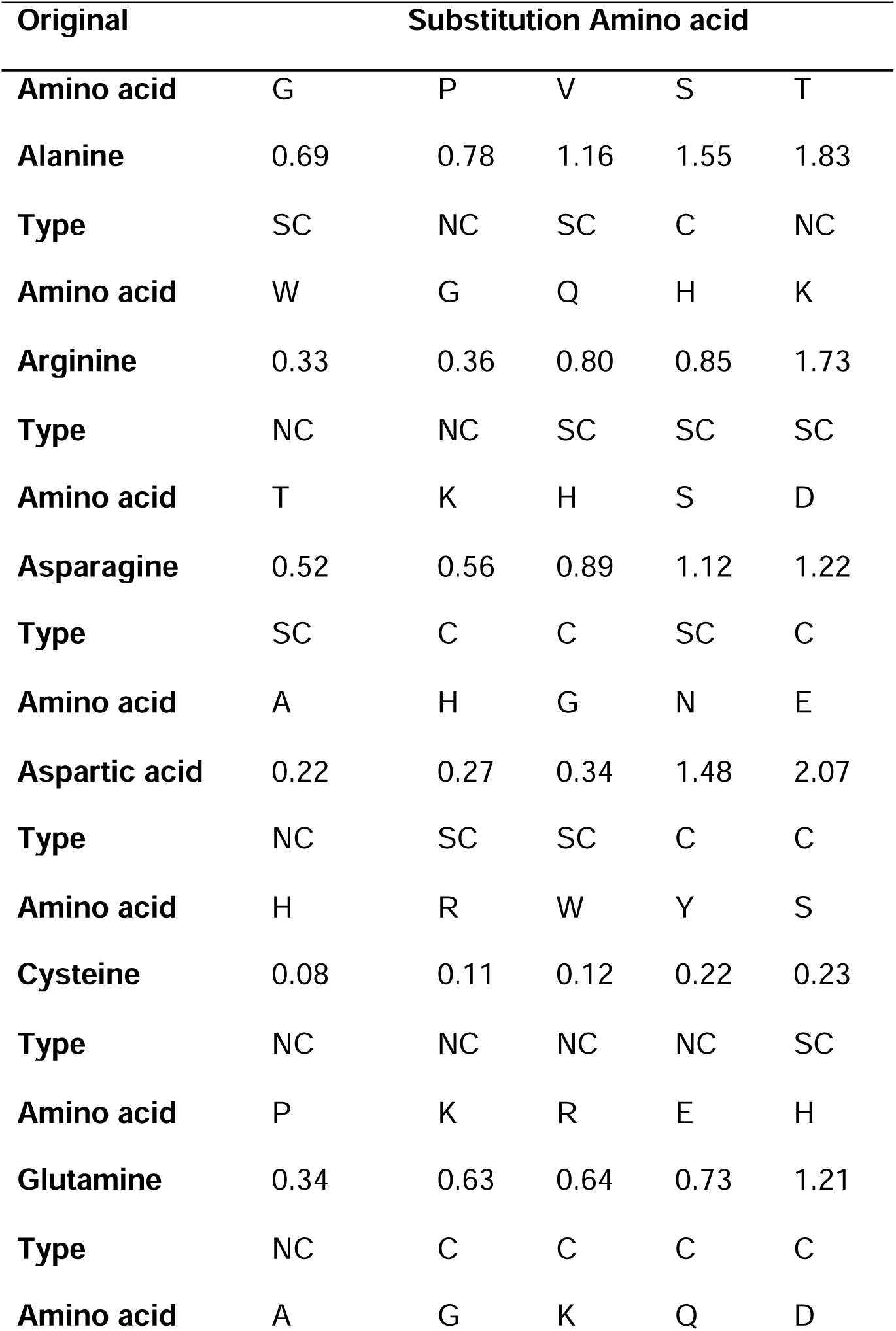
Randomly selected amino acids to assess type of substitutions from Maximum Likelihood Estimate of Substitution Matrix. C: Conservative substitutions, SC-Semi-conservative substitutions, NC: Non-conservative substitutions.

**S5 Table.** Tajima Relative Test. Neutrality substitution rate was calculated for PSTOL1 using the taxas: A (*T. aestivum* 3DL-*PSTOL1*) and B (ANC28927 *O. sativa* PSTOL1) with sequence C (A. tauschii *PSTOL1*). For Lr10Kinase, A (ABB84341 *T. aestivum*) and B (XP 021311718 S. bicolor Lr10RLPK2.3X1), C (XP 020184941 *A.tauschii* Lr10RLPK2.4X3). For *PSTOL1*/*LrK10* Kinase genes A (*T. aestivum* 3DL-*PSTOL1*) and B (XP 014758757 *B. distachyon* RLPK), with sequence C (XP 020166263 *A. tauschii* Lr10RLPK2.1). The χ^2^ test statistic was 13.09 (P = 0.00030 with 1 degree of freedom). P-value less than 0.05 is often used to reject the null hypothesis of equal rates between lineages. The analysis carried out with MEGA 7, the analysis consists of 3 amino acid sequences.

| Configuration | *PSTOL1* | *Lr10Kinase* | *PSTOL1/Lr10Kinase* |
| --- | --- | --- | --- |
| Identical sites in all three sequences | 198 | 164 | 250 |
| Divergent sites in all three sequences | 7 | 44 | 9 |
| Unique differences in Sequence A | 2 | 107 | 10 |
| Unique differences in Sequence B | 81 | 11 | 34 |
| Unique differences in Sequence C | 1 | 5 | 22 |
| P-value | 0.0000 | 0.0000 | 0.0003 |
| $\chi^2$ | 75.19 | 78.10 | 13.09 |
| Degree of freedom | 1 | 1 | 1 |

**S6 Table.** SPCP-structural and physico-chemical properties. NAA-Number of amino acids, AH-Alpha helix, ES-Extended strand, RC-Random coil, pI-Theoretical Isoelectric focusing point, NCR-Negatively charged residues (Aspartic acid and Glutamic acid), PCR-Positively charged residues (Arginine and Lysine), ID-Instability Index, AI-Aliphatic index. P3DL-*PSTOL1*:3DL, *LrK10*-Leaf Rust Resistance Kinase, *LrK10A*-Leaf Rust Resistance Kinase *PSTOL1* Aligned Region, P3AL-*PSTOL1*:3AL, P5AS-*PSTOL1*:5AS. PAt-*PSTOL1*-*A. tauchii*, *RK2*-*T. aestivum Receptor like kinase 2,* RK2A-*T. aestivum Receptor like kinase 2* aligned region.

| SPCP | P3DL | LrK10 | LrK10A | P3AL | P5AS | PAt | RK2 | RK2A |
| --- | --- | --- | --- | --- | --- | --- | --- | --- |
| **NAA** | 339 | 636 | 350 | 377 | 289 | 339 | 634 | 350 |
| **AH** | 138 | 227 | 170 | 158 | 120 | 131 | 223 | 162 |
| **ES** | 59 | 104 | 39 | 63 | 51 | 67 | 108 | 43 |
| **RC** | 142 | 305 | 141 | 156 | 118 | 141 | 303 | 145 |
| **pI** | 8.52 | 6.34 | 8.23 | 8.67 | 6.24 | 8.67 | 8.53 | 9.07 |
| **NCR** | 38 | 69 | 38 | 38 | 33 | 37 | 61 | 35 |
| **PCR** | 42 | 64 | 40 | 44 | 31 | 42 | 69 | 42 |
| **ID** | 36.42 | 44.57 | 36.61 | 36.35 | 30.31 | 36.29 | 42.53 | 33.90 |
| **AI** | 86.25 | 87.22 | 91.89 | 89.97 | 81.97 | 87.96 | 87.93 | 92.14 |

